# Inference of Locus-Specific Population Mixtures From Linked Genome-Wide Allele Frequencies

**DOI:** 10.1101/2023.11.06.565831

**Authors:** Carlos S. Reyna-Blanco, Madleina Caduff, Marco Galimberti, Christoph Leuenberger, Daniel Wegmann

## Abstract

Admixture between populations and species is common in nature. Since the influx of new genetic material might be either facilitated or hindered by selection, variation in mixture proportions along the genome is expected in organisms undergoing recombination. Various graph-based models have been developed to better understand these evolutionary dynamics of population splits and mixtures. However, current models assume a single mixture rates for the entire genome and do not explicitly account for linkage. Here, we introduce `TreeSwirl`, a novel method for inferring branch lengths and locus-specific mixture proportions by using genome-wide allele frequency data, assuming that the admixture graph is known or has been inferred. `TreeSwirl` builds upon `TreeMix` that uses Gaussian processes to estimate the presence of gene flow between diverged populations. However, in contrast to `TreeMix`, our model infers locus-specific mixture proportions employing a Hidden Markov Model that accounts for linkage. Through simulated data, we demonstrate that `TreeSwirl` can accurately estimate locus-specific mixture proportions and handle complex demographic scenarios. It also outperforms related D- and f-statistics in terms of accuracy and sensitivity to detect introgressed loci.

## 2 Introduction

Gene flow, the exchange of genetic material between populations or different species (Slatkin, 1985a), can occur through various mechanisms, such as migration, admixture, hybridization, cross-fertilization, or even by the dispersal of diaspores and pollinators (Barton and Hewitt, 1985; Ellstrand et al., 2003; Tung and Barreiro, 2017; Burgarella et al., 2019). This exchange may play a significant role in the maintenance of genetic variation, but also in the adaptation to multiple ecological niches (Anderson, 1949; Slatkin, 1985b, 1987; Rieseberg and Wendel, 1993; Barton, 2001). At sufficient levels, gene flow can lead to homogenization of populations, particularly in the face of opposing genetic drift (Ellstrand, 2014). Gene flow might also increase genetic variation at a much higher rate than mutation (Grant and Grant, 1994) and impact the process of speciation by becoming a primary source of genetic diversity and adaptive novelty for a population (Ellstrand et al., 2003; Abbott et al., 2013). Several genetic analyses have shown that gene flow, both ancient and present, is a common phenomenon in nature (Grant and Grant, 1992; Mallet, 2005; Patterson et al., 2006; Wegmann and Excoffier, 2010; Tung and Barreiro, 2017; Marchi et al., 2022), and a bifurcating tree, representing population or species historical relationships, fails to account for it (Kulathinal et al., 2009; Reich et al., 2009; Sousa et al., 2009; Green et al., 2010; Durand et al., 2011; Reich et al., 2012). This led to the development of methods that use allele-frequency data and graph-based models to infer population splits and test for the presence of gene flow between divergent populations or species (Pickrell and Pritchard, 2012; Patterson et al., 2012; Yang et al., 2012; Eaton and Ree, 2013; Lipson et al., 2013, 2014; Martin et al., 2013; Kozak et al., 2021), which, for instance, confidently settled the long-standing question whether gene flow occurred between modern humans and archaic hominins. However, these methods assume a genome-wide gene flow rate per migration edge, which is unrealistic in the presence of selection. In theory, the effective gene flow may vary significantly along the genome because of selection and genetic drift (Yamamichi and Innan, 2012), making it essential to quantify these variations to better understand the dynamics that lead to introgression (Racimo et al., 2015, 2017; Suarez-Gonzalez et al., 2018; Sankararaman, 2020).

Introgression is a lasting consequence of gene flow that leads to the assimilation of variants into the local gene pool through repeated back-crossing, resulting in their permanent inclusion (Anderson and Hubricht, 1938). When introgressed loci increase the fitness of the recipient population, this is known as “adaptive introgression”. Unlike neutral introgression, which can be lost over time due to drift, adaptive introgression is sustained by selection and can eventually lead to fixation (Zhang et al., 2021). But selection may also prevent introgression if it reduces fitness (Christe et al., 2016). While introgressed loci may be identified through explicit demographic modelling (Luqman et al., 2021), the classic way is via population genetic summary statistics. Patterson’s D, for example, has been estimated in sliding windows along the genome to identify introgressed loci (Dasmahapatra et al., 2012; Kronforst et al., 2013; Smith and Kronforst, 2013; Rheindt et al., 2014; Fontaine et al., 2015). Since it was originally intended for genome-wide analysis (Martin et al., 2015), more suitable related statistics have been used for analyzing specific short genomic regions, such as *f*_*d*_, *f*_*dM*_, and *d*_*f*_ (Martin et al., 2015; Malinsky et al., 2015; Pfeifer and Kapan, 2019; Malinsky et al., 2021). There are other statistics, for instance, S* and its variants that use linkage disequilibrium information to detect long introgressed haplotypes (Plagnol and Wall, 2006; Wall et al., 2009; Vernot and Akey, 2014; Vernot et al., 2016; Browning et al., 2018) or ArchIE that combines diverse summary statistics to detect introgressed haplotypes without a reference (Durvasula and Sankararaman, 2019, 2020). However, outlier scans based on such statistics are likely to ignore valuable information present in the full data, do not model linkage explicitly or require an arbitrary choice of large window-size and outliers identification. To overcome these constraints, probabilistic frameworks such as Hidden Markov Models (HMMs) (Rabiner and Juang, 1986; Prüfer et al., 2014; Seguin-Orlando et al., 2014; Skov et al., 2018; Steinrücken et al., 2018), and Conditional Random Fields (CRF) (Sankararaman et al., 2014) have been applied to infer the ancestry state of each site. These methods are extensions of models that infer local ancestry from genotyping data (Tang et al., 2006; Price et al., 2009; Wegmann et al., 2011; Lawson et al., 2012; Maples et al., 2013) and while explicitly accounting for demographic history and linkage, they rely on phased and training sequence data, unadmixed or archaic reference, and detailed demographic models. As a consequence, such approaches are not easily applicable to non-model species for which only limited data and knowledge is available.

To complement these methods, we here propose a model that makes use of Gaussian processes to infer locus-specific mixture proportions. Gaussian processes have a rather long history to model allele frequency differences between populations (Cavalli-Sforza and Edwards, 1967; Felsenstein, 1981), but have recently seen a surge in applications due to the development of the popular tool `TreeMix` (Pickrell and Pritchard, 2012). Our method, `TreeSwirl`, explicitly takes an admixture graph (e.g. inferred by TreeMix) and genome-wide allele frequencies to infer locus-specific mixture proportions. To account for linkage, we make use of a Hidden Markov Model (HMM), wherein the hidden states are represented by the proportion of the mixture at a particular site and the observed data is represented by the sampled allele frequencies. To evaluate the performance of our method against other tools, we simulated data using various demographic models. We estimated the mixture proportions with `TreeSwirl` and computed related D- and F-statistics using D-suite Dinvestigate (Malinsky et al., 2021). Our findings revealed that `TreeSwirl` surpasses the summary statistics estimates in detecting the simulated signal of introgression under different scenarios, although at an additional computational cost. Furthermore, by applying `TreeSwirl` to real data cases, we successfully identified candidate genomic regions where migration rates fluctuate and may be subject to selection.

## 3 Materials and Methods

### 3.1 The Model

Consider a set of populations *m* = 1, 2, …, *M* that are linked by a graph 𝒢 which represents their population history in terms of population splits and migration events. Consider as well a series of diploid, bi-allelic loci *l* = 1, 2, …, *L*, where the total number of loci *L* might constitute, for instance, consecutive SNPs along the genome. At each locus *l*, a total number of ***N*** _*l*_ = (*N*_*l*1_, …, *N*_*lM*_) alleles have been observed across the *M* populations, of which ***n***_*l*_ = (*n*_*l*1_, …, *n*_*lM*_) were derived and the remaining ancestral (or otherwise polarized). To model sampled allele counts ***n***_*l*_|***N***_*l*_ we distinguish two processes: the first models the distribution of the vector of the actual but unknown population frequencies ***y***_*l*_ = (*y*_*l*1_, …, *y*_*lM*_) ^*′*^ given the graph 𝒢, and the second the distribution of the sampled allele counts ***n***_*l*_|***N*** _*l*_ given ***y***_*l*_ (Fig 1A).

**Figure 1:**
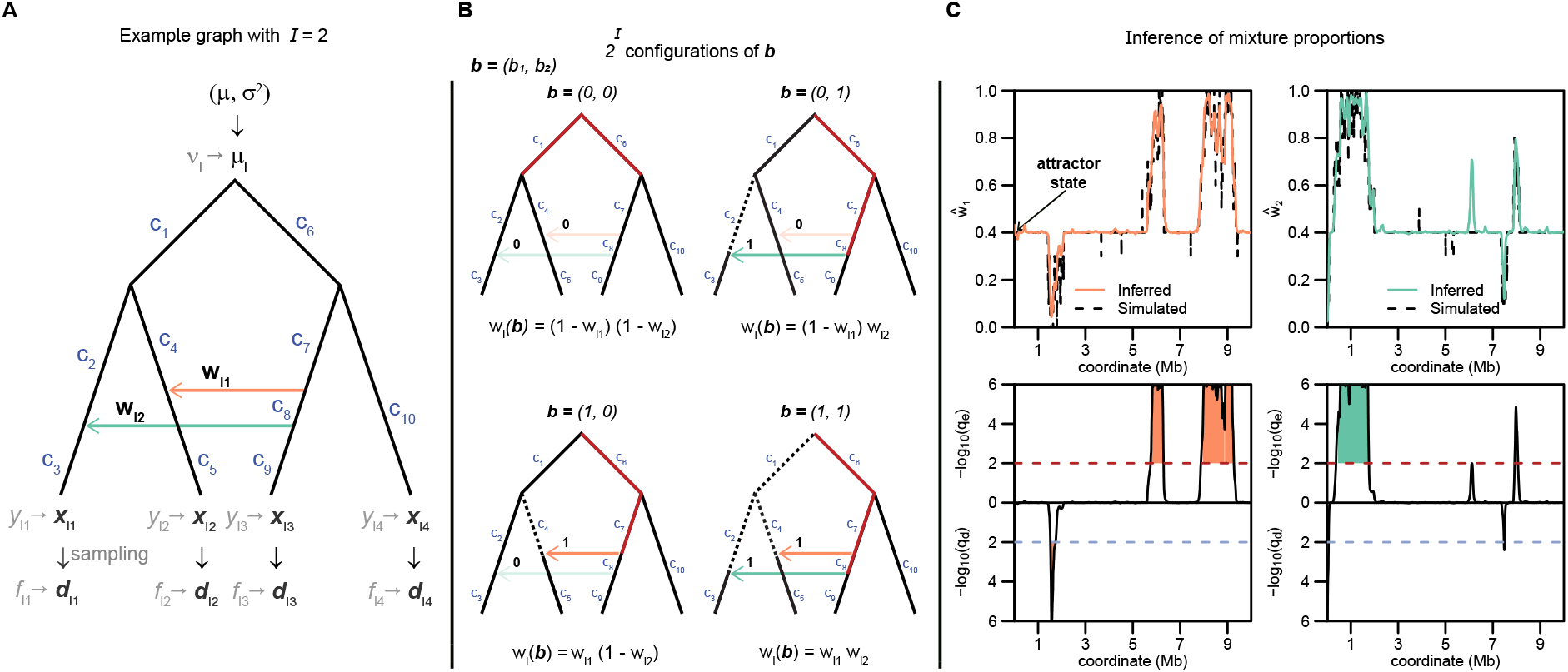
Inference example. A: admixture graph with two migration edges marked in different colors. Parameters of interest are shown on the graph (root prior and branch lengths) as well as the untransformed and transformed ancient, sampling and population allele frequency variables. B: All possible configurations of **b** for two migration events when they are open or closed. C: Example of inference under our `TreeSwirl` model for each migration event. The top panel shows the posterior mean mixture proportions *ŵ*_*l*_ compared to simulated estimates and the bottom panel shows the identified candidate regions under possible selection, where the false discovery rates (FDR) for excess (*q*_*e*_) and dearth (*q*_*d*_) introgression was determined for each locus as explained in the “Inference” section.

#### 3.1.1 Evolution along the graph 𝒢

We assume, as in Pickrell and Pritchard (2012), that the change in allele frequencies from the root to the tips of 𝒢 is modeled as a Brownian motion (BM) process. For each locus *l*, the BM process starts at the root of 𝒢 at a value of allele frequency which we denote by *ν*_*l*_. It proceeds along the branches of 𝒢 and finally gives rise to the above-mentioned random vector ***y***_*l*_ at the leaves of G. The probability of ***y***_*l*_ is given by the multivariate normal density

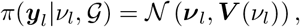

where ***ν***_*l*_ = (*ν*_*l*_, …, *ν*_*l*_)^*′*^ is the mean vector and ***V***(*ν*_*l*_) is the variance-covariance matrix corresponding to the BM on𝒢. For the construction of ***V***(*ν*_*l*_), which depends on the topology of 𝒢, the lengths of the branches in 𝒢 and the migration rates, we follow Pickrell and Pritchard (2012). We set

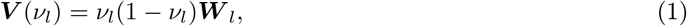

where ***W*** _*l*_ only depends on the graph topology, the branch lengths and the migration rates.

However, it was long recognized that BM with constant variance is not adequately describing allele frequency changes, especially close to boundaries and various transformations to alleviate the problem have been proposed (Felsenstein, 1981). Here we will consider the transformation

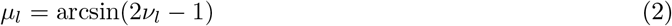

from the interval [0, 1] onto [−*π/*2, *π/*2]. This has the advantage that all factors of *ν*_*l*_(1 − *ν*_*l*_) in front of the variance matrices will be canceled. We thus replace (eq. 1) by

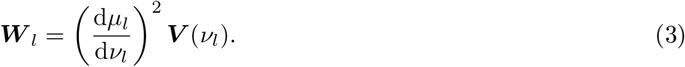

Let ***x***_*l*_ = (*x*_*l*1_, …, *x*_*lM*_), *x*_*lm*_ = arcsin(2*y*_*lm*_ − 1) denote the transformed population allele frequencies. The distribution of ***x***_*l*_ thus follows the multivariate normal density

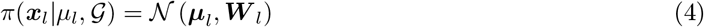

with ***µ***_*l*_ = (*µ*_*l*_, …, *µ*_*l*_) = *µ*_*l*_**1**.

The matrix ***W*** _*l*_ is constructed as follows. Let 𝒢 be a rooted population graph with *K* oriented branches *k* = 1, …, *K* of length *c*_*k*_, ***c*** = (*c*_1_, …, *c*_*K*_)^*′*^; the orientation of the branches points in direction of the leaves. We assume that the graph also contains *I* oriented migration edges *τ*_*i*_, *i* = 1, …, *I*, to which we assign no branch length. The migration edges should be placed such that there are no cycles in the graph. However, we allow for bi-directional migration edges, which, to avoid cycles, we model effectively as two migration edges, of which the starting point of one precedes the end point of the other by an infinitesimally small branch.

We now consider paths leading from the root of the graph to a leaf taking some of the migration edges (open edges) and leaving others out (closed edges). More precisely, let

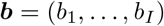

be a binary vector indicating a certain configuration of open and closed migration edges: a bit *b*_*i*_ = 1 indicates that the migration edge *τ*_*i*_ is open and *b*_*i*_ = 0 that the migration edge *τ*_*i*_ is closed (Fig 1B). We denote by *w*_*li*_ the migration rate, i.e. the probability of edge *τ*_*i*_ to be open, and thus we assign to the configuration ***b*** the probability

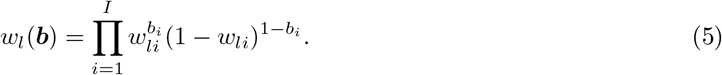

Now, for a given configuration ***b***, pick a population (leaf) *m* and a branch *k*. There is at most one path leading from the root to the population *m* and taking exactly the open migration edges according to ***b***. If, moreover, this path contains the branch *k*, we set the indicator function *I*_*mk*_(***b***) equal to 1. Otherwise we set *I*_*mk*_(***b***) = 0.

Using this notation, we can now define the *M* × *M*-matrices ***J*** _*lk*_ for each branch *k* element-wise by

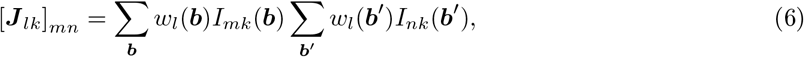

where each sum runs over all the 2^*I*^ possible configurations of ***b*** and ***b***^*′*^, respectively. Each matrix ***J*** _*lk*_ thus reflects the probabilities that branch *k* was common for any pair of leaves.

The matrix ***W*** _*l*_, after all, is given by

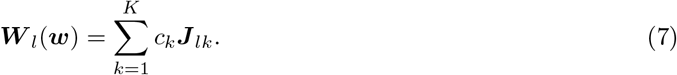

This construction of the variance matrix ***W*** _*l*_(***w***_*l*_) is a generalized reformulation of an argument given in Pickrell and Pritchard (2012).

To unclutter the notation, we will use ***W*** _*l*_ = ***W*** _*l*_(***w***_*l*_) in the rest of this article and thus not indicate its dependence on the migration rates ***w***_*l*_ = (*w*_*l*1_, …, *w*_*lI*_).

#### 3.1.2 Sampling

We assume that the observed allele counts *n*_*lm*_ at locus *l* and population *m* follow a binomial distribution with parameters *N*_*lm*_ and *y*_*lm*_, where *y*_*lm*_ is the true allele frequency in population *m*. By independence of the samples, we have

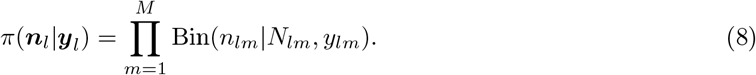

If the sample sizes are sufficiently large, we can approximate this distribution by a multivariate normal density. Let **f**_*l*_ = (*f*_*l*1_, …, *f*_*lM*_) with *f*_*lm*_ = *n*_*lm*_*/N*_*lm*_ denote the observed allele frequencies at locus *l*, which are approximately normally distributed with with mean ***y***_*l*_ and a diagonal variance-covariance matrix:

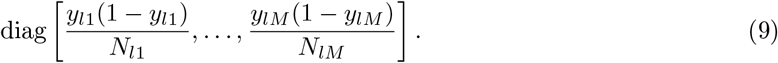

The transformed observed allele frequencies **d**_*l*_ = (*d*_*l*1_, …, *d*_*lM*_) with *d*_*lm*_ = arcsin(2*f*_*lm*_ − 1), are then approximated by the multivariate normal density

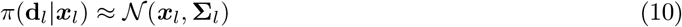

with

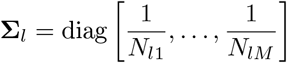

because the factors *y*_*l*1_(1 − *y*_*l*1_) are transformed away from the variance-covariance matrix (eq. 9) similar to (eq. 3).

#### 3.1.3 Full likelihood for one locus

Given the ancestral frequency *µ*_*l*_, we obtain the likelihood by combining (eq. 4) and (eq. 10) and integrating out:

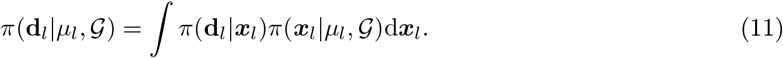

Using well-known formulae for linear systems (see Thm. 4.4.1 in Murphy (2012)) we obtain for the likelihood (eq. 11) the following approximation:

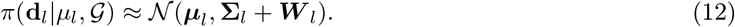

We now set a normal prior on *µ*_*l*_, namely we assume that

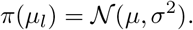

Again from Thm. 4.4.1 in Murphy (2012) we conclude that

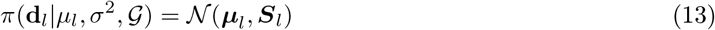

with

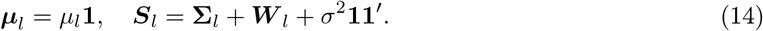

Explicitly

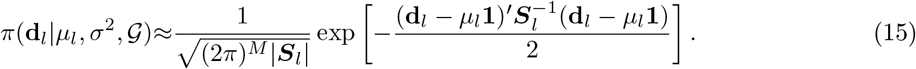

We note that the above model accounts for variation in sample size among loci, but assumes that at least one sample was observed per locus and population.

### 3.2 Hidden Markov Model

We develop a Hidden Markov Model (HMM) for multiple loci *l* = 1, …, *L* with varying migration rates for each of the *I* migration edges of graph G. We assume that the locus and specific migration rates *w*_*li*_ take values out of a small set of discrete numbers between 0 and 1:

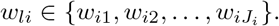

We thus have *J*_1_ · *J*_2_ · … · *J*_*I*_ possible combinations and these combinations will constitute the hidden states of our Markov model. We denote the hidden state at locus *l* by *z*_*l*_. Each state *z*_*l*_ corresponds to a multiindex

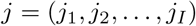

that defines the migration values 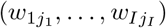 of the migration edges. Thus, knowing the state *z*_*l*_ is tantamount to knowing the combination of migration rates at the given site which in turn determines the matrix ***W*** in eq. (eq. 7) via (eq. 5) and (eq. 6).

To account for linkage between loci, we assume that the locus-specific transition matrix 𝕡 (*z*_*l*_ = *j*^*′*^|*z*_*l−*1_ = *j*) is based on physical or genetic distances *δ*_*l*_ between loci. We assume independence of the transition probabilities of the different migration edges:

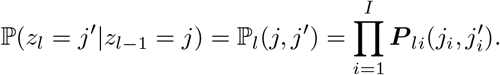

Each one of the factors in this product is an element of a ladder-type Markov matrix ***P*** _*li*_ which is defined via a transition rate matrix *κ*_*i*_**Λ**_*i*_:

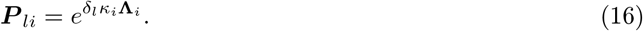

Here, *κ*_*i*_ is a positive scaling parameter pertaining to migration edge *i*, the distances *δ*_*l*_ are known constants corresponding to the linking distances. Further, the *J*_*i*_ × *J*_*i*_-matrices **Λ**_*i*_ reflect a transition model for infinitesimal steps.

We consider two transition models: The first is a standard ladder-type model in which transitions are only allowed to neighboring states and at equal rates:

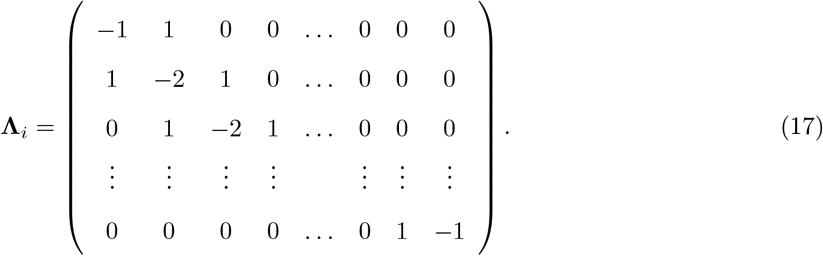

Under this transition matrix, only *κ*_*i*_ is inferred, for which we maximize the Q-function using a linear search.

The second is also a ladder-type model, but which includes an attractor state 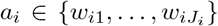 reflecting the background migration rate. Similar to Galimberti et al. (2020), we use two parameters to describe the fraction of loci deviating from the attractor state and the degree of that deviation, but chose a slightly different parametrization. Specifically, and given the two parameters *ϕ*_*i*_ and *ζ*_*i*_, we have

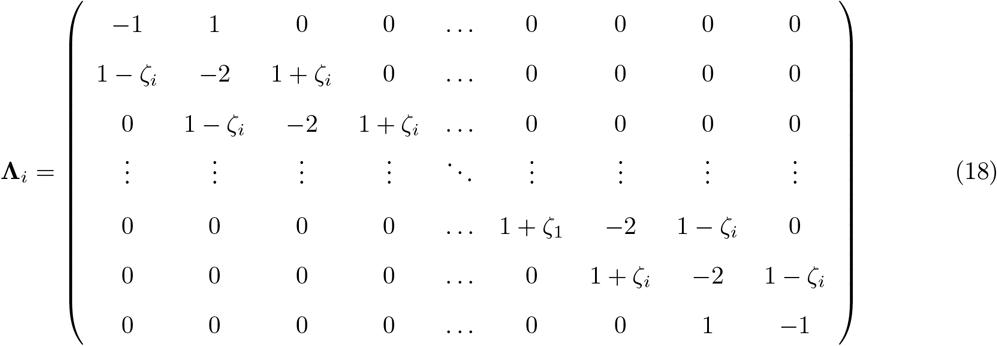

with the attractor row given by

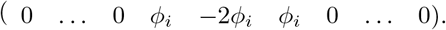

Note that the *κ*_*i*_, *ϕ*_*i*_ and *ζ*_*i*_ all must be strictly positive. However, we limit *ϕ*_*i*_ and *ζ*_*i*_ to the range (0,1] to ensure that the stationary probability of the attractor state *a*_*i*_ is higher than for any other state.

Finally, the emission probabilities are generated via the marginal likelihood (eq. 15):

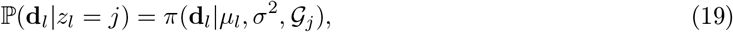

where 𝒢_*j*_ denotes the population graph with migration rates according to the state *z*_*l*_ = *j* and *µ*_*l*_ is the root state at site *l*.

### 3.3 Inference

We developed an empirical Bayes inference scheme for the hidden states under the assumption that the topology of the admixture graph is either known or was previously obtained. Specifically, we first infer both the emission and transition probabilities using the Baum-Welch algorithm (Baum et al., 1970) and then posterior state probabilities under the inferred parameters. As detailed in the Supplementary Information (see section “Inference”), the Baum-Welch algorithm requires numerical optimization in each iteration. While the parameter of the root prior *µ* can be optimized analytically, we resort to Newton-Raphson optimization (Nocedal and Wright, 2006; Lange, 2010) for the root prior *σ*_2_ and for parameters of the population graph (i.e. the branch lengths ***c***) and to Nelder-Mead optimization (Nelder and Mead, 1965) for the parameters regarding the transition matrices with attractors (i.e. the *κ*_*i*_, *ϕ*_*i*_, *ζ*_*i*_) or a linear search for transition matrices with no attractors (i.e. the *κ*_*i*_).

The Baum-Welch algorithm may be sensitive to initial conditions. We obtain initial estimates of all parameter values as follows (see Supplementary Information for more details):

1. We use the observed variance-covariance matrix of the transformed observed frequencies as an initial guess of the variance covariance matrix ***W***.
2. To account for variation in ***W*** among loci, we refine this initial estimate using a Gaussian Mixture Model (GMM) under which the transformed observed frequencies are modeled by one of *r* = 1, …, *R* multivariate Gaussian distributions with variance-covariances matrices ***W*** _*r*_ but shared root priors *µ* and *σ*_2_. This model assumes no constraints regarding the structure of the ***W*** _*r*_ and can be optimized with an Expectation-Maximization (EM) algorithm with analytic updates. The hidden states ***s*** = (*s*_1_, …, *s*_*L*_) with *s*_*l*_ ∈ {1, …, *R*} are assumed to follow a Categorical distribution ***s*** ∼ Cat(***π***) with ***π*** = (*π*_1_, …, *π*_*R*_). We impose the constraint that each *π*_*r*_ is larger than some threshold, which we set to 0.2 by default. If this constraint is not met upon convergence, we decrease *R* = *R* − 1 and repeat this step.
3. We next use a Nelder-Mead algorithm to coerce the inferred variance-covariance matrices ***W*** _1_, …, ***W*** _*R*_ onto the population graph. Specifically, we seek to find the set of branch lengths ***c*** and partition-specific migration rate ***w***_*r*_ that best explain the previously learned variance-covariance matrices using the weighted Residuals Sum of Squares. To mitigate convergence problems, we repeat the Nelder-Mead *C* times and use different initial values each time. We set *C* = 1000 by default.
4. We then run a Baum-Welch algorithm to infer the branch lengths ***c***, the root prior *µ* and *σ*_2_ and the transition parameters *κ*_*i*_ for the attractor-free, ladder-type transition matrix (eq. 17). Upon convergence, we identify the top *T* states with the highest average posterior probabilities across all loci, and consider each of these states as a potential attractor. We set *T* = 5 by default.
5. To identify the best attractor, we then run individual Baum-Welch optimization for each potential attractor, thereby identifying the best values for the branch lengths ***c***, the root prior *µ* and *σ*_2_ and the transition parameters *κ*_*i*_, *ϕ*_*i*_ and *ζ*_*i*_ for the full transition matrix (eq. 18). The final result, as relevant to the user, are the results with the transition matrix that resulted in the highest likelihood, which may either be the ladder-type matrix or one with a specific attractor state.

Once maximum likelihood estimates for the branch lengths ***c***, the transition parameters *κ*_*i*_, *ϕ*_*i*_, *ζ*_*i*_ and *a*_*i*_ as well as the root prior *µ* and *σ*_2_ are obtained, we infer state posterior probabilities *P* (*z*_*l*_|**d, *θ***) given the full data **d** and the learned parameters collectively denoted by ***θ***, see Fig 1C. We further determined the posterior mean migration rates as

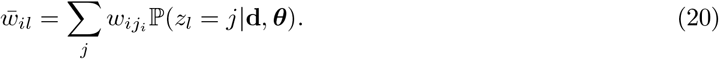

To identify regions exhibiting either excess or dearth introgression compared to the genome-wide average, and are hence candidate regions to have experienced selection, we summarized these posterior probabilities as

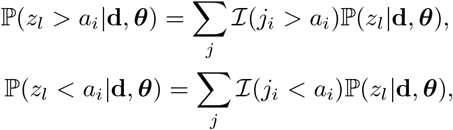

where ℐ (·) denotes the indicator function. We then determined for each locus *l* the false discovery rates (FDR) for excess (*q*_*e*_(*l*)) and dearth (*q*_*d*_(*l*)) introgression as

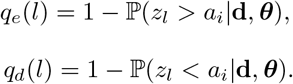

### 3.4 Implementation

We implemented the proposed inference scheme as a user-friendly C++ program `TreeSwirl`, which is available, along with documentation, through a git repository at bitbucket.org/wegmannlab/treeswirl. Our implementation makes heavy use of the HMM framework of the statistical library `stattools` available at bitbucket.org/wegmannlab/stattools.

To streamline computations, we employ a straightforward clustering method to reduce the number of sampling size variance matrices **Σ**_*l*_ that need to be considered:

1. We sort the vector of sample sizes according to the frequency of each occurrence.
2. To cluster, we identify the pair of vectors with the least occurrences and compute their weighted average.
3. We retain the weighted vector of sample sizes, remove the pair, and update the occurrence count as the sum of the deleted pair counts.
4. We repeat steps 1 through 3 until the desired number of **Σ**_*l*_ is obtained.

Given a limited number *u* of such matrices and given that we use a finite number of discrete migration rates, there exist also an only finite number of matrices ***S***_*l*_ that can be pre-computed in each Baum-Welch iteration to speed up the forward-backward pass through the HMM.

### 3.5 Simulations

#### 3.5.1 Simulations under model assumptions

To demonstrate the power and limitations of our methods with respect to different demographic scenarios, we simulated data under the `TreeSwirl` model for population graphs of four populations under four different topologies, visualized in Figure 2. In the first two cases, we designed a tree with a single migration event starting ancestral to Population 2 and directed to either Population 3 (Topology 1) or ancestral to Population 3 (Topology 2). The remaining two cases had two migration events, either bidirectional migration ancestral to the Populations 2 and 3, or two unrelated events from ancestral of Population 2 and 4 to ancestral of Populations 3 and 1, respectively. For each tree, we simulated a single chromosome and considered both a scenario with a constant, genome-wide migration rate, as well as a scenario where the migration rate varied. For all cases, we ran 20 replicates with *N* = 100, *µ* = −0.5, *σ* = 0.3 and *J* = 21.

**Figure 2:**
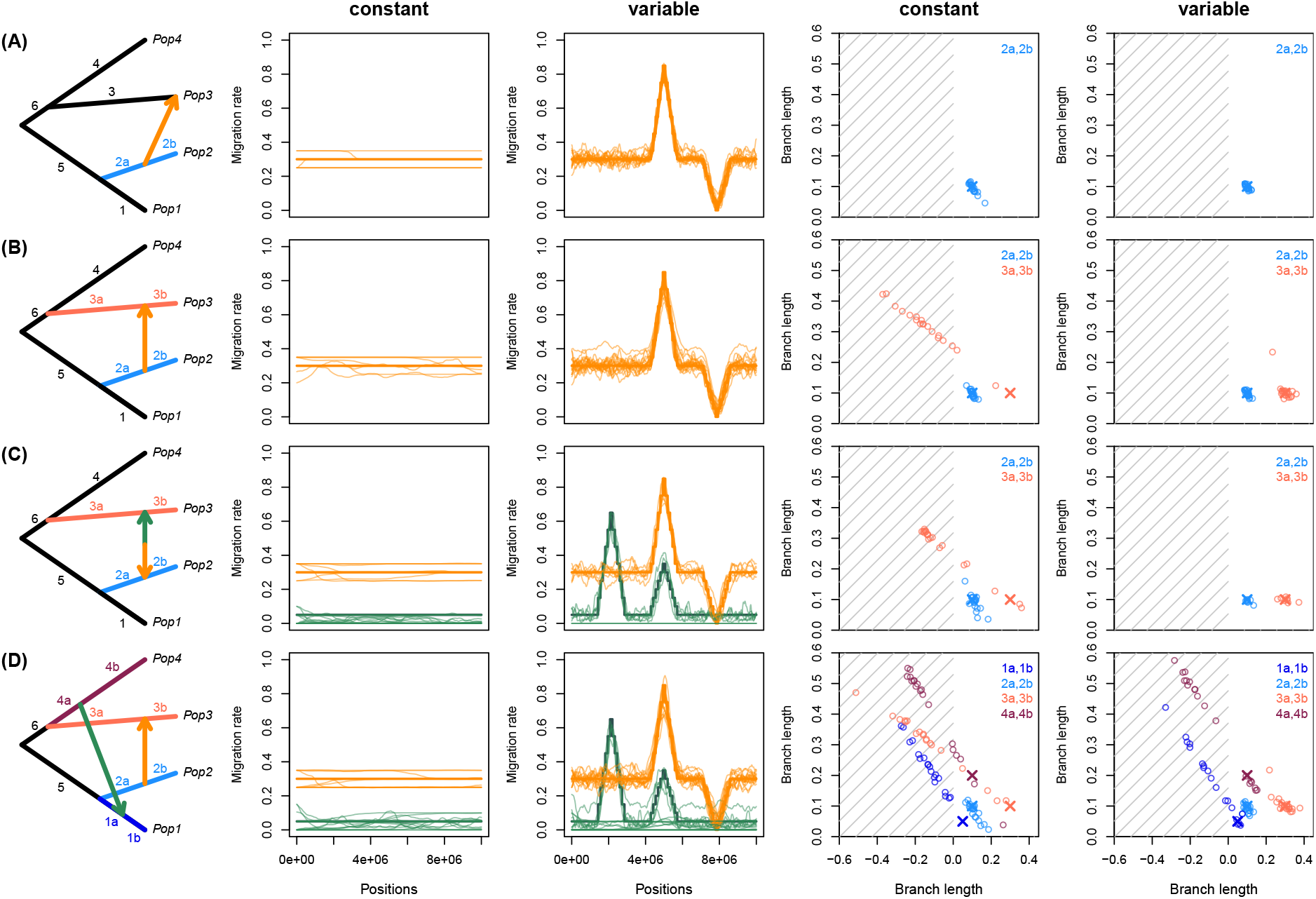
Performance of `TreeSwirl` as assessed by simulations under the `TreeSwirl` model. Shown are, for each of four cases of migration scenarios, the population graph (first column), estimates of locus-specific migration rates (posterior means, second and third column) and estimates of selected branch lengths (fourth and fifth column) for a case with variable (third and fifth column) and for a case without (second and fourth column) variation in migration rates along the simulated chromosome. Colors correspond to the migration edges and branch lengths of the same color in the population graph. Simulated migration rates are shown as a thick line and the estimates as a thin line per replicate. For branch lengths the simulated values are shown as a cross and estimates as open circles.

#### 3.5.2 fastsimcoal2

To compare `TreeSwirl` to competing methods, we used `fastsimcoal2` (Excoffier et al., 2021) to simulate genomic data under five different demographic scenarios only consisting of population splits and admixture pulses (but no population growth or continuous migration, Figure 3). We maintained a constant effective population size of *N*_*e*_ = 10, 000 and used a sample size of *N* = 100 for each population in all cases.

**Figure 3:**
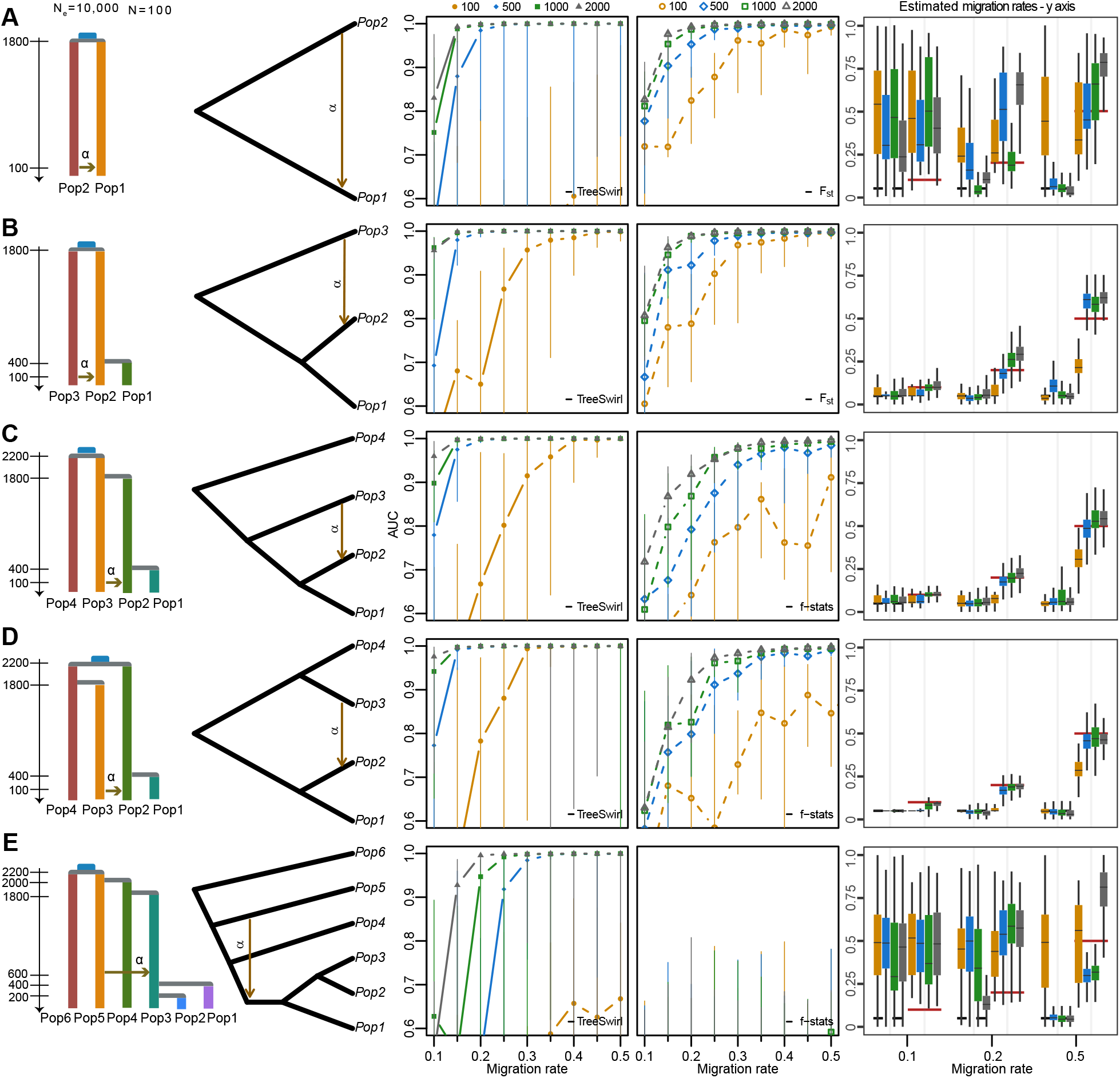
Performance of `TreeSwirl` and *f*_4_-stats methods to measure the amount of introgression under different demographic histories with an background migration rate *α*_*b*_ = 0.05. First column: simulated demographic histories. Second column: schematic of the topology models. Third column: AUC measures for `TreeSwirl` as a function of the migration rate of blocks with elevated migration rates. Fourth column: AUC measure for the combination of summary statistic (*F*_*ST*_, *D* − or *f* -related), window size and population subset (in case E only) that led to the highest AUC. Symbols indicate the size of the blocks with elevated migration rates compared to the background rate.

To simulate variation in admixture pulses along chromosomes, we composed each chromosome of seven blocks, each containing many independent DNA segments of length 1000 bp, fully-linked (i.e. within-locus recombination rate of 0.0), a mutation rate of 1e-8, and a transition rate of 0.33. Odd-numbered blocks reflected the neutral genomic background, each containing *n*_*b*_ = 3, 500 loci and an admixture pulse of *α*_*b*_ = 0.05. Conversely, even-numbered blocks reflected loci under selection. While all three selected blocks shared the same parameters in one simulation, we varied their number of DNA segments *n*_*s*_ and migration rates *α*_*s*_ across different simulations.

We generated 10 replicates for each parameter combination and used a custom script to transform the generated output files into standard VCF files and concatenating the seven blocks corresponding to a single chromosome. We then applied a minimum allele frequency filter of maf = 0.05 with VCFtools (Danecek et al., 2011), resulting in a variable but rather low (0-3) number of loci per DNA segment and thus resulted in data similar to what would be obtained after LD-pruning.

The filtered VCFs served as input for estimating sliding window *F*_*st*_ for simulated data only consisting of two or three populations as well as for running `D-suite Dinvestigate` (Malinsky et al., 2021) with varying window sizes *s* = (10, 50, 100, 150, 200, 250, 300, 350, 400, 450, 500), a sliding locus of 1, and the true trio and corresponding outgroup for demographic scenarios with more than three populations. Concurrently, we executed `TreeSwirl` using the same filtered data and the expected tree topology.

We employed a receiver operating characteristic (ROC) curve analysis to assess the area under the curve (AUC), which summarizes the performance of the method in distinguishing introgression from the “neutral state” *ix*. For the ROC analysis, we used the estimated mean posteriors obtained from `TreeSwirl`, along with the computed values of *F*_*st*_, Patterson’s D, *f*_*d*_, *f*_*dM*_, and *d*_*f*_. For each comparison, we used the statistics and window-size that resulted in the best AUC.

### 3.6 Real data application

We re-analyzed whole genome sequencing from a recent study that identified putatively adaptive introgression from domestic goat into Alpine ibex (Münger et al., 2023). We restricted our analysis to species with at least four individuals: modern Alpine ibex (29 individuals), the Iberian ibex (four individuals), Bezoar ibex (six individuals) and domestic goats (16 individuals). We further focused on Chromosomes 23 on which significant introgression was previously reported for gene regions with immune-related genes such as MHC (Grossen et al., 2014; Münger et al., 2023), and included the flanking chromosomes 21, 22, 24 and 25 for comparison. We used a VCF file provided by the authors. This file was previously filtered to a minimum minor allele frequency (MAF) of 0.05 and a maximal missingness of 0.9 across all samples studied in Münger et al. (2023), and was further thinned to have a minimal distance of 100 bp between consecutive SNPs (see Münger et al., 2023, for details). The admixture graph used was derived from the demographic model inferred by Münger et al. (2023), but we modelled two migration events starting ancestral to domestic goat and directed to Alpine ibex and one to Iberian ibex, respectively, since hybridization between domestic goats and Iberian ibex has also been reported (Cardoso et al., 2021). We used the physical positions of markers provided in the VCF file and set *J* = 21.

## 4 Results

### 4.1 Simulations under the TreeSwirl model

We conducted simulations under the above model to investigate the power of `TreeSwirl` to infer branch lengths and locus-specific migration rates. We focused on a tree of four populations and considered several cases of migration events of increasing complexity (Fig 2) and simulated a case with and without variation in migration rates along the simulated chromosome.

The first case contains a single migration event from a population ancestral to Population 2 to Population 3. In this case, the most challenging branch lengths to estimate are those leading to and from the source of the migration edge (branches 2a and 2b in Fig 2A) since only their sum is relevant for the variance across loci in Population 2. However, additional information about their lengths comes from the covariance between populations 2 and 3, and as a result their lengths as well as locus-specific migration rates are well estimated both when using a constant or a variable migration rate (Fig 2A).

The second case is similar to the first, except that the migration edge now ends ancestral to Population 3, leading to two pairs of branches that are challenging to infer: branches 2a and 2b and branches 3a and 3b (marked in blue and pink in Fig 2B, respectively). As in the first case, information about the lengths of branches 2a and 2b stems from the variance of Population 2 and the covariance between populations 2 and 3. In contrast, information about the lengths of branches 3a and 3b only stems from the variance of Population 3, and they are thus non-identifiable in the case of constant migration with some branch lengths often inferred to be negative (Fig 2B). This non-identifiability issue has been previously observed by Pickrell and Pritchard (2012), and `TreeMix` consequently only allows for migration edges that end at tips or the end of branches. As shown in Fig 2B, however, these branch lengths do become identifiable if loci vary in their migration rates and if that variation is explicitly modelled.

And as shown in Fig 2C, variation in migration rates renders all branch lengths identifiably also in the case of bi-directional migration, which led to an accurate inference of locus-specific migration rates in all simulated replicates. While variation in migration rates renders all branch lengths identifiable also in the case of two migration events involving all four populations, our inference framework fails to identify the correct solutions in about 2/3 of all replicates (Fig 2D). Closer inspection reveals that this mostly affects the branch lengths and migration rates related to the migration edge that is more ancient and has both lower average and less variability in migration rates. We note that these cases often result in negative branch lengths, which we found to be a good indicator that the population graph used has too many degrees of freedom for the data analyzed.

### 4.2 Comparison to related D- and f-statistic methods

We generated coalescent simulations from five demographic histories of population splits and mixtures with `fastsimcoal2`. We used an effective population size of *N*_*e*_ = 10, 000, a sample size of *N* = 100 and a shared common ancestor for all populations dating back approximately 2,000 generations (Fig 3, first column). Each simulated chromosome involved seven genomic blocks: odd blocks were 3,500 loci long and simulated using a background migration rate of *α*_*b*_ = 0.05, while even blocks were shorter and had elevated migration rates. We used a range of block lengths and elevated migration rates to evaluate the power of `TreeSwirl` and commonly used summary statistics to identify regions with elevated migration rates, namely *F*_*st*_ and f-statistics. We calculated these statistics using `D-suite Dinvestigate` (Malinsky et al., 2021) for various window sizes and, in case of more than four populations, for all population combinations possible. To render our comparison conservative, wen then always kept the summary statistics, window size and population combination resulting in highest power. The second column of Fig 3 shows the population graph used for `TreeSwirl`.

As shown in the third and fourth column of Fig 3, `TreeSwirl` has overall higher power to identified regions with elevated migration rates (candidates for adaptive introgression) than competing summary statistics, with a much lower false-positive rate, particularly if the background and elevated migration rates were rather similar. Importantly, `TreeSwirl` also had considerably power in identifying loci with elevated migration rates for models with two- and three-taxon topologies. This presents a significant advantage over *f*_4_-stat methods, which are constrained to four-taxon configurations and require a pre-defined outgroup.

The most difficult scenario was the case of only two populations, in which `TreeSwirl` needs to estimate migration rates along four branch lengths. In the case of no variation in migration rates, the information in the data corresponds to only two variances and one covariance term, thus considerably less than the degrees of freedom in the model. By only simulating two different migration rates in the genome, as done here, there is still not enough information to render branch lengths and migration rates identifiable as evidence by the failure of `TreeSwirl` to accurately infer the migration strength. Nonetheless, `TreeSwirl` was more accurate in identifying the loci with elevated migration rates than *F*_*st*_ in all cases except when the blocks with elevated migration were very short (100 loci) or when the elevated migration rate was very close to the background rate (Fig 3A).

As more populations are included, the number of variances and covariances increases faster than the number of branch lengths (*M* (*M* + 1)*/*2 vs. 2*M* − 1 for a strictly bifurcating graph without migration), rendering model parameters easier to infer (but see above for how migration edges complicate things). Consequently, `TreeSwirl` also inferred migration rates much more accurately for simulations conducted with more populations (Fig 3B-D), although it remained difficult in the case of very short blocks of elevated migration (100 loci).

As illustrated in Fig 3E, also models with many populations may be challenging to infer. In that scenario, the migration edge is more ancient and the populations Pop1 and Pop2 do not help to resolve the difficulty in inferring the length of the branches leading to and from the source and destination of the migration edge. The scenario thus corresponds to that explored above in Fig 2B, which is only identifiable with sufficient variation in migration rates. Interestingly, and despite its rather low power, `TreeSwirl` still outperformed all summary statistics tested in identifying the loci with elevated migration rates.

### 4.3 Introgression from domestic goats into Alpine ibex

Alpine ibex are a wild goat species native to the European alps. It suffered from a near-extinction two centuries ago, but has since recovered thanks to re-population efforts. It is known that Alpine ibex occasionally hybridize with domestic goats and adaptive introgression has been reported for several genomic regions, particularly for a region on Chromosome 23 that contains immune-related genes such as MHC (Grossen et al., 2014; Münger et al., 2023). Here we used `TreeSwirl` to re-analyze whole-genome sequencing data of Alpine ibex, Iberian ibex, Bezoar ibex and domestic goat to quantify locus-specific rates of introgression from domestic goat into Alpine and Iberian ibex along chromosomes 21-25. As shown in Figure 4, we generally inferred very low migration rates from domestic goat into Alpine ibex with an inferred attractor state at zero. Nonetheless, we identified a clear signal of introgression on Chromosomes 23 between 20-24Mb, corresponding well to the region previously reported (Münger et al., 2023) and containing MHC. We also identified and additional, smaller region of introgression on Chromosome 21.

**Figure 4:**
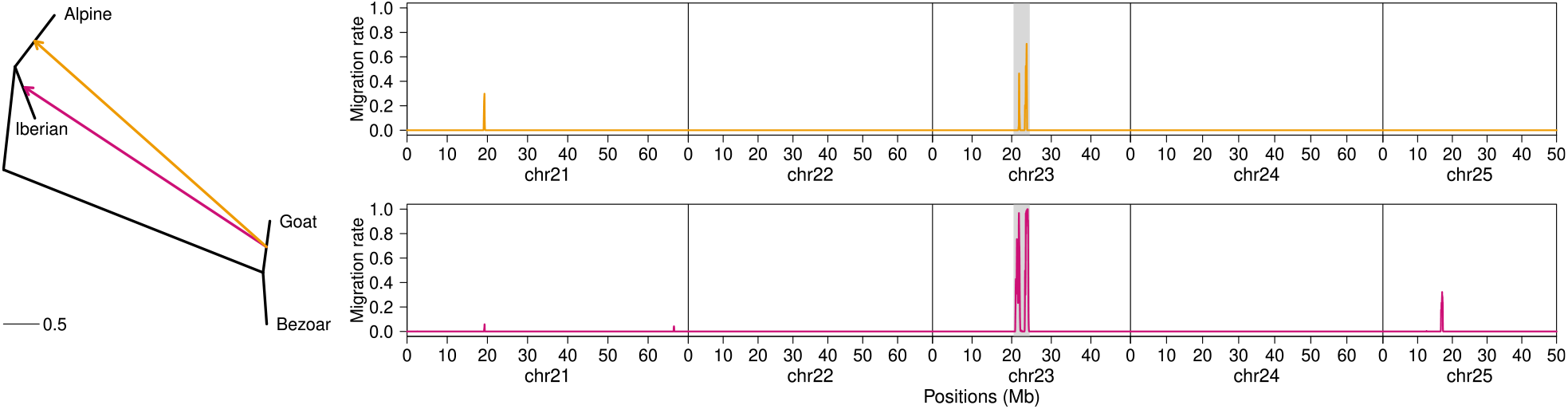
Inference of introgressed loci in Alpine and Iberian ibex. Shown is the inferred population graph along with inferred migration rates (posterior means) along chromosomes 21-25 for Alpine ibex (top, orange) and Iberian ibex (bottom, pink). The region reported by Münger et al. (2023) on Chromosome 23 is marked with a gray background.

We also inferred rather low overall migration rates from domestic goat into Iberian ibex, again with the attractor state estimated at zero. Interestingly, the region on Chromosome 23 inferred as introgressed in Alpine ibex was also inferred as strongly introgressed in Iberian ibex, this time along an additional, smaller region on Chromosome 25. Since the two ibex species likely diverged prior to the domestication of goats, the signal on Chromosome 23 is thus indicative of independent adaptive introgression in both ibex species. But we note that given the frequency dependent selection likely acting on the MHC region, the signal may also be indicative of shared ancient polymorphisms among all goat species, although the high migration rates we inferred (close to 1.0) suggest this scenario to be less likely.

### 4.4 Runtime considerations

The computational performance of `TreeSwirl` is influenced by multiple factors, such as the number of discrete states *J*, the number of matrices **Σ**, and the total number of sites and migration edges. Computation times scale exponentially with the number of migration edges, quadratically with the number of states *J* and linearly with the number of loci. Computations may be efficiently distributed across multiple computer nodes by dividing the genome into independent segments, such as individual chromosomes or chromosome arms. This approach is valid because linkage does not persist across chromosome boundaries and is typically weak across the centromere.

## 5 Discussion

One approach to infer historical relationships among populations is to model allele frequency changes along a phylogenetic tree as a Gaussian process (Cavalli-Sforza and Edwards, 1967; Felsenstein, 1981). This rather old concept was recently revived by extending the model to a graph with migration edges and by providing a user-friendly tool `TreeMix` to infer parameters under such a graph (Pickrell and Pritchard, 2012). However, the `TreeMix` model assumes migration rates to be constant along the genome, an assumption that may not hold in the face of selection or strong genetic drift. Indeed, theory predicts variation in the rate of effective gene flow along the genome (Harrison, 1993), in which local barriers to gene flow are anticipated to emerge from the random accumulation of Dobzhansky-Muller incompatibilities, both under models of secondary contact after isolation (Barton and Gale, 1993) as well as under models of continuous gene flow during speciation (Wu, 2001). In the case of gene flow between highly divergent gene pools, selection is likely to act as the primary driving force for variation in effective gene flow along the genome, with rates of introgression being particularly low in genomic regions involved in adaptation, so called islands of speciation, but potentially much higher in regions free from the selection pressure (Dasmahapatra et al., 2012).

In light of these considerations, we here present `TreeSwirl`, an extension of the model described in Pickrell and Pritchard (2012) that allows for mixture proportions to vary along the genome in an auto-correlated way that reflects the effect of linkage. We evaluated the performance of our model to identify such variation in comparison to existing methods related to *D*- and *f* -statistics, such as *F*_*st*_, Patterson’s *D* (Patterson et al., 2012), *f*_*d*_ (Martin et al., 2015), *f*_*dM*_ (Malinsky et al., 2015), and *d*_*f*_ (Pfeifer and Kapan, 2019), which have been frequently applied to identify signatures of introgression using arbitrary genomic window sizes. As we show using extensive simulations, our method had superior accuracy and sensitivity in detecting retrogressed loci under a wide range of demographic histories characterized by single admixture pulses.

The approach presented here also addresses numerous constraints inherent to the use of related *D*- and *f* -statistics. First, these summary statistics are limited to bifurcating four-population topologies. In cases involving graphs of five or more populations, the simplest option is to subsample a section of the graph in the appropriate configuration, as done in `Dsuite` (Malinsky et al., 2021) used here. In cases involving two- or three-population topologies, one would need to resort to *F*_*ST*_ -based metrics. In contrast, the method presented here is not constraint by topology, working well with any number of populations and also under topologies that include polytomies.

Second, our HMM-based approach to model linkage eliminates the need to specify window sizes. Instead, the parameters governing auto-correlation are directly inferred from the data along with introgression rates. In our simulations, the choice of window sizes, as well as the choice of the specific statistics to use, had a big impact on power. To ensure a fair comparison between methods, we thus tested all available summary statistics for a wide range of window sizes and only report the results of the combination of summary statistics and window size that was optimal for each individual case. In applications to real data, however, such explorations are not possible, likely leading to an even larger difference in power between `TreeSwirl` and these summary statistics.

Third, `TreeSwirl` supports graphs with multiple migration edges for which introgression rates are learned simultaneously. However, it is important to note that the performance of `TreeSwirl` is likely dependent on the quality of the tree topology used as input and may not perform well if the tree topology is poorly resolved or incorrect. Similarly, we caution that `TreeSwirl` just as `TreeMix` assumes that population allele frequencies can be explained through a population graph with splits, drift along branches and potential mixtures. While this approach has been proved useful in many cases, it is easy to think of scenarios that violate these assumptions. One such scenario consists of populations that have diverged long enough to hardly share any polymorphisms. This leads to negative covariance between populations (the presence of a mutation in one population implies its absence in the others), which can not be captured by a population graph. While experimenting with such cases showed `TreeSwirl` to be remarkably robust, it should not be confused with phylogenetic approaches.

Other scenarios that violate the underlying model assumptions are cases of extensive continuous gene flow as well as cases of population structure not well captured with a graph. Such scenarios were shown to lead to spurious inference of introgression using other methods (e.g. Eriksson and Manica, 2014; Lawson et al., 2018; Tournebize and Chikhi, 2023), and while we have not explored these scenarios in detail here, we are convinced they can lead to spurious inference by both `TreeSwirl` and `TreeMix` and appeal to users to carefully assess model assumptions of these any other tools they use. Nonetheless, and as we showed through both simulations and a data application, the model introduced here is powerful to identify introgressed loci under many scenarios and will thereby contribute to a better understanding of the role of introgression in evolution, which is expected to act as a major driver in adaptation to ongoing global changes (Suarez-Gonzalez et al., 2018).

## Data and code availability

The authors affirm that all data required to validate the conclusions of this article are either included within the article itself or accessible through the indicated repositories. The source code for `TreeSwirl` can be found in the following Git repository: bitbucket.org/wegmannlab/treeswirl2, which also contains a user manual. This study did not generate any new data.

## Acknowledgments

We thank Christine Grossen for sharing the ibex data and two anonymous reviewer for their helpful comments on an earlier version of this manuscript. This work was supported by Swiss National Science Foundation grants 31003A_173062 and 310030_200420 to DW.

## Author contributions

DW conceived the idea; DW, CL and CSRB developed the model; CSRB and MC implemented the method in collaboration with MG; CSRB and MC conducted all simulations and data analyses; CSRB, MC and DW led the writing of the manuscript. All authors contributed critically to the draft and gave final approval for publication.

## References

Abbott, R., Albach, D., Ansell, S., Arntzen, J. W., Baird, S. J., Bierne, N., Boughman, J., Brelsford, A., Buerkle, C. A., Buggs, R., Butlin, R. K., Dieckmann, U., Eroukhmanoff, F., Grill, A., Cahan, S. H., Hermansen, J. S., Hewitt, G., Hudson, A. G., Jiggins, C., Jones, J., Keller, B., Marczewski, T., Mallet, J., Martinez-Rodriguez, P., Möst, M., Mullen, S., Nichols, R., Nolte, A. W., Parisod, C., Pfennig, K., Rice, A. M., Ritchie, M. G., Seifert, B., Smadja, C. M., Stelkens, R., Szymura, J. M., Väinölä, R., Wolf, J. B., and Zinner, D. (2013). Hybridization and speciation. Journal of Evolutionary Biology, 26(2):229–246.

Anderson, E. (1949). Introgressive hybridization. J. Wiley, New York.

Anderson, E. and Hubricht, L. (1938). Hybridization in Tradescantia. III. The Evidence for Introgressive Hybridization. American Journal of Botany, 25(6):396.

Barton, N. and Gale, K. (1993). Genetic analysis of hybrid zones. In Harrison, R. G., editor, Hybrid Zones and the Evolutionary Process, pages 13–45. Oxford University Press.

Barton, N. H. (2001). The role of hybridization in evolution. Molecular Ecology, 10(3):551–568.

Barton, N. H. and Hewitt, G. M. (1985). Analysis of Hybrid Zones. Annual Review of Ecology, Evolution, and Systematics, pages 113–148.

Baum, L. E., Petrie, T., Soules, G., and Weiss, N. (1970). A Maximization Technique Occurring in the Statistical Analysis of Probabilistic Functions of Markov Chains. The Annals of Mathematical Statistics, 41(1):164–171.

Browning, S. R., Browning, B. L., Zhou, Y., Tucci, S., and Akey, J. M. (2018). Analysis of Human Sequence Data Reveals Two Pulses of Archaic Denisovan Admixture. Cell, 173(1):53–61.e9.

Burgarella, C., Barnaud, A., Kane, N. A., Jankowski, F., Scarcelli, N., Billot, C., Vigouroux, Y., and Berthouly-Salazar, C. (2019). Adaptive introgression: An untapped evolutionary mechanism for crop adaptation. Frontiers in Plant Science, 10:4.

Cardoso, T. F., Luigi-Sierra, M. G., Castelló, A., Cabrera, B., Noce, A., Mármol-Sánchez, E., García-González, R., Fernández-Arias, A., Alabart, J. L., López-Olvera, J. R., Mentaberre, G., Granados-Torres, J. E., Cardells-Peris, J., Molina, A., Sànchez, A., Clop, A., and Amills, M. (2021). Assessing the levels of intraspecific admixture and interspecific hybridization in iberian wild goats (capra pyrenaica). Evolutionary Applications, 14(11):2618–2634.

Cavalli-Sforza, L. L. and Edwards, A. W. (1967). Phylogenetic analysis. Models and estimation procedures. American Journal of Human Genetics, 19(3 Pt 1):233.

Christe, C., Stölting, K. N., Bresadola, L., Fussi, B., Heinze, B., Wegmann, D., and Lexer, C. (2016). Selection against recombinant hybrids maintains reproductive isolation in hybridizing populus species despite f1 fertility and recurrent gene flow. Molecular Ecology, 25(11):2482–2498.

Danecek, P., Auton, A., Abecasis, G., Albers, C. A., Banks, E., DePristo, M. A., Handsaker, R. E., Lunter, G., Marth, G. T., Sherry, S. T., McVean, G., and Durbin, R. (2011). The variant call format and VCFtools. Bioinformatics, 27(15):2156–2158.

Dasmahapatra, K. K., Walters, J. R., Briscoe, A. D., Davey, J. W., Whibley, A., Nadeau, N. J., Zimin, A. V., Salazar, C., Ferguson, L. C., Martin, S. H., Lewis, J. J., Adler, S., Ahn, S. J., Baker, D. A., Baxter, S. W., Chamberlain, N. L., Ritika, C., Counterman, B. A., Dalmay, T., Gilbert, L. E., Gordon, K., Heckel, D. G., Hines, H. M., Hoff, K. J., Holland, P. W., Jacquin-Joly, E., Jiggins, F. M., Jones, R. T., Kapan, D. D., Kersey, P., Lamas, G., Lawson, D., Mapleson, D., Maroja, L. S., Martin, A., Moxon, S., Palmer, W. J., Papa, R., Papanicolaou, A., ick Pauchet, Y., Ray, D. A., Rosser, N., Salzberg, S. L., Supple, M. A., Surridge, A., Tenger-Trolander, A., Vogel, H., Wilkinson, P. A., Wilson, D., Yorke, J. A., Yuan, F., Balmuth, A. L., Eland, C., Gharbi, K., Thomson, M., Gibbs, R. A., Han, Y., Jayaseelan, J. C., Kovar, C., Mathew, T., Muzny, D. M., Ongeri, F., Pu, L. L., Qu, J., Thornton, R. L., Worley, K. C., Wu, Y. Q., Linares, M., Blaxter, M. L., Ffrench-Constant, R. H., Joron, M., Kronforst, M. R., Mullen, S. P., Reed, R. D., Scherer, S. E., Richards, S., Mallet, J., Mc Millan, W. O., and Jiggins, C. D. (2012). Butterfly genome reveals promiscuous exchange of mimicry adaptations among species. Nature, 487(7405):94–98.

Durand, E. Y., Patterson, N., Reich, D., and Slatkin, M. (2011). Testing for ancient admixture between closely related populations. Molecular Biology and Evolution, 28(8):2239–2252.

Durvasula, A. and Sankararaman, S. (2019). A statistical model for reference-free inference of archaic local ancestry. PLOS Genetics, 15(5):e1008175.

Durvasula, A. and Sankararaman, S. (2020). Recovering signals of ghost archaic introgression in African populations. Science Advances, 6(7).

Eaton, D. A. and Ree, R. H. (2013). Inferring Phylogeny and Introgression using RADseq Data: An Example from Flowering Plants (Pedicularis: Orobanchaceae). Systematic Biology, 62(5):689–706.

Ellstrand, N. C. (2014). Is gene flow the most important evolutionary force in plants? American Journal of Botany, 101(5):737–753.

Ellstrand, N. C., Barrett, S. C., Linington, S., Stephenson, A. G., and Comai, L. (2003). Current knowledge of gene flow in plants: implications for transgene flow. Philosophical Transactions of the Royal Society of London. Series B: Biological Sciences, 358(1434):1163–1170.

Eriksson, A. and Manica, A. (2014). The Doubly Conditioned Frequency Spectrum Does Not Distinguish between Ancient Population Structure and Hybridization. Molecular Biology and Evolution, 31(6):1618–1621.

Excoffier, L., Marchi, N., Marques, D. A., Matthey-Doret, R., Gouy, A., and Sousa, V. C. (2021). fastsimcoal2: demographic inference under complex evolutionary scenarios. Bioinformatics, 37(24):4882–4885.

Felsenstein, J. (1981). Evolutionary trees from gene frequencies and quantitative characters: Finding maximum likelihood estimates. Evolution; international journal of organic evolution, 35(6):1229–1242.

Fontaine, M. C., Pease, J. B., Steele, A., Waterhouse, R. M., Neafsey, D. E., Sharakhov, I. V., Jiang, X., Hall, A. B., Catteruccia, F., Kakani, E., Mitchell, S. N., Wu, Y. C., Smith, H. A., Rebecca Love, R., Lawniczak, M. K., Slotman, M. A., Emrich, S. J., Hahn, M. W., and Besansky, N. J. (2015). Extensive introgression in a malaria vector species complex revealed by phylogenomics. Science, 347(6217):1258524.

Galimberti, M., Leuenberger, C., Wolf, B., Szilágyi, S. M., Foll, M., and Wegmann, D. (2020). Detecting Selection from Linked Sites Using an F-Model. Genetics, 216(4):1205–1215.

Grant, P. R. and Grant, B. R. (1992). Hybridization of Bird Species. Science, 256(5054):193–197.

Grant, P. R. and Grant, B. R. (1994). Phenotypic and genetic effects of hybridization in Darwin’s finches. Evolution, 48(2):297–316.

Green, R. E., Krause, J., Briggs, A. W., Maricic, T., Stenzel, U., Kircher, M., Patterson, N., Li, H., Zhai, W., Fritz, M. H. Y., Hansen, N. F., Durand, E. Y., Malaspinas, A. S., Jensen, J. D., Marques-Bonet, T., Alkan, C., Prüfer, K., Meyer, M., Burbano, H. A., Good, J. M., Schultz, R., Aximu-Petri, A., Butthof, A., Höber, B., Höffner, B., Siegemund, M., Weihmann, A., Nusbaum, C., Lander, E. S., Russ, C., Novod, N., Affourtit, J., Egholm, M., Verna, C., Rudan, P., Brajkovic, D., Kucan, Ž., Gušic, I., Doronichev, V. B., Golovanova, L. V., Lalueza-Fox, C., De La Rasilla, M., Fortea, J., Rosas, A., Schmitz, R. W., Johnson, P. L., Eichler, E. E., Falush, D., Birney, E., Mullikin, J. C., Slatkin, M., Nielsen, R., Kelso, J., Lachmann, M., Reich, D., and Pääbo, S. (2010). A draft sequence of the neandertal genome. Science, 328(5979):710–722.

Grossen, C., Keller, L., Biebach, I., Consortium, T. I. G. G., and Croll, D. (2014). Introgression from domestic goat generated variation at the major histocompatibility complex of alpine ibex. PLOS Genetics, 10(6):1–16.

Harrison, R. G. (1993). Hybrid Zones and the Evolutionary Process. Oxford University Press.

Kozak, K. M., Joron, M., McMillan, W. O., and Jiggins, C. D. (2021). Rampant Genome-Wide Admixture across the Heliconius Radiation. Genome Biology and Evolution, 13(7).

Kronforst, M. R., Hansen, M. E., Crawford, N. G., Gallant, J. R., Zhang, W., Kulathinal, R. J., Kapan, D. D., and Mullen, S. P. (2013). Hybridization Reveals the Evolving Genomic Architecture of Speciation. Cell reports, 5(3):666.

Kulathinal, R. J., Stevison, L. S., and Noor, M. A. (2009). The Genomics of Speciation in Drosophila: Diversity, Divergence, and Introgression Estimated Using Low-Coverage Genome Sequencing. PLOS Genetics, 5(7):e1000550.

Lange, K. (2010). Numerical Analysis for Statisticians. Statistics and Computing. Springer New York, New York, NY.

Lawson, D. J., Hellenthal, G., Myers, S., and Falush, D. (2012). Inference of Population Structure using Dense Haplotype Data. PLOS Genetics, 8(1):e1002453.

Lawson, D. J., Van Dorp, L., and Falush, D. (2018). A tutorial on how not to over-interpret structure and admixture bar plots. Nature communications, 9(1):3258.

Lipson, M., Loh, P. R., Levin, A., Reich, D., Patterson, N., and Berger, B. (2013). Efficient Moment-Based Inference of Admixture Parameters and Sources of Gene Flow. Molecular Biology and Evolution, 30(8):1788–1802.

Lipson, M., Loh, P. R., Patterson, N., Moorjani, P., Ko, Y. C., Stoneking, M., Berger, B., and Reich, D. (2014). Reconstructing Austronesian population history in Island Southeast Asia. Nature Communications 2014 5:1, 5(1):1–7.

Luqman, H., Widmer, A., Fior, S., and Wegmann, D. (2021). Identifying loci under selection via explicit demographic models. Molecular Ecology Resources, 21(8):2719–2737.

Malinsky, M., Challis, R. J., Tyers, A. M., Schiffels, S., Terai, Y., Ngatunga, B. P., Miska, E. A., Durbin, R., Genner, M. J., and Turner, G. F. (2015). Genomic islands of speciation separate cichlid ecomorphs in an East African crater lake. Science, 350(6267):1493–1498.

Malinsky, M., Matschiner, M., and Svardal, H. (2021). Dsuite - Fast D-statistics and related admixture evidence from VCF files. Molecular Ecology Resources, 21(2):584–595.

Mallet, J. (2005). Hybridization as an invasion of the genome. Trends in Ecology & Evolution, 20(5):229– 237.

Maples, B. K., Gravel, S., Kenny, E. E., and Bustamante, C. D. (2013). RFMix: A Discriminative Modeling Approach for Rapid and Robust Local-Ancestry Inference. The American Journal of Human Genetics, 93(2):278–288.

Marchi, N., Winkelbach, L., Schulz, I., Brami, M., Hofmanová, Z., Blöcher, J., Reyna-Blanco, C. S., Diekmann, Y., Thiéry, A., Kapopoulou, A., Link, V., Piuz, V., Kreutzer, S., Figarska, S. M., Ganiatsou, E., Pukaj, A., Struck, T. J., Gutenkunst, R. N., Karul, N., Gerritsen, F., Pechtl, J., Peters, J., Zeeb-Lanz, A., Lenneis, E., Teschler-Nicola, M., Triantaphyllou, S., Stefanović, S., Papageorgopoulou, C., Wegmann, D., Burger, J., and Excoffier, L. (2022). The genomic origins of the world’s first farmers. Cell, 185(11):1842–1859.e18.

Martin, S. H., Dasmahapatra, K. K., Nadeau, N. J., Salazar, C., Walters, J. R., Simpson, F., Blaxter, M., Manica, A., Mallet, J., and Jiggins, C. D. (2013). Genome-wide evidence for speciation with gene flow in Heliconius butterflies. Genome research, 23(11):1817–1828.

Martin, S. H., Davey, J. W., and Jiggins, C. D. (2015). Evaluating the Use of ABBA–BABA Statistics to Locate Introgressed Loci. Molecular Biology and Evolution, 32(1):244–257.

Münger, X., Robin, M., Dalén, L., and Grossen, C. (2023). Facilitated introgression from domestic goat into alpine ibex at immune loci. bioRxiv.

Murphy, K. P. (2012). Machine learning: a probabilistic perspective.

Nelder, J. A. and Mead, R. (1965). A simplex method for function minimization. The Computer Journal, 7(4):308–313.

Nocedal, J. and Wright, S. (2006). Numerical Optimization. Springer Science & Business Media.

Patterson, N., Moorjani, P., Luo, Y., Mallick, S., Rohland, N., Zhan, Y., Genschoreck, T., Webster, T., and Reich, D. (2012). Ancient admixture in human history. Genetics, 192(3):1065–1093.

Patterson, N., Richter, D. J., Gnerre, S., Lander, E. S., and Reich, D. (2006). Genetic evidence for complex speciation of humans and chimpanzees. Nature 2006 441:7097, 441(7097):1103–1108.

Pfeifer, B. and Kapan, D. D. (2019). Estimates of introgression as a function of pairwise distances. BMC Bioinformatics, 20(1):1–11.

Pickrell, J. K. and Pritchard, J. K. (2012). Inference of Population Splits and Mixtures from Genome-Wide Allele Frequency Data. PLOS Genetics, 8(11):e1002967.

Plagnol, V. and Wall, J. D. (2006). Possible ancestral structure in human populations. Plos Genetics, 2(7):e105–e105.

Price, A. L., Helgason, A., Palsson, S., Stefansson, H., St. Clair, D., Andreassen, O. A., Reich, D., Kong, A., and Stefansson, K. (2009). The Impact of Divergence Time on the Nature of Population Structure: An Example from Iceland. PLOS Genetics, 5(6):e1000505.

Prüfer, K., Racimo, F., Patterson, N., Jay, F., Sankararaman, S., Sawyer, S., Heinze, A., Renaud, G., Sudmant, P. H., De Filippo, C., Li, H., Mallick, S., Dannemann, M., Fu, Q., Kircher, M., Kuhlwilm, M., Lachmann, M., Meyer, M., Ongyerth, M., Siebauer, M., Theunert, C., Tandon, A., Moorjani, P., Pickrell, J., Mullikin, J. C., Vohr, S. H., Green, R. E., Hellmann, I., Johnson, P. L., Blanche, H., Cann, H., Kitzman, J. O., Shendure, J., Eichler, E. E., Lein, E. S., Bakken, T. E., Golovanova, L. V., Doronichev, V. B., Shunkov, M. V., Derevianko, A. P., Viola, B., Slatkin, M., Reich, D., Kelso, J., and Pääbo, S. (2014). The complete genome sequence of a Neanderthal from the Altai Mountains. Nature 2013 505:7481, 505(7481):43–49.

Rabiner, L. R. and Juang, B. H. (1986). An Introduction to Hidden Markov Models. IEEE ASSP Magazine, 3(1):4–16.

Racimo, F., Marnetto, D., and Huerta-Sánchez, E. (2017). Signatures of Archaic Adaptive Introgression in Present-Day Human Populations. Molecular Biology and Evolution, 34(2):296–317.

Racimo, F., Sankararaman, S., Nielsen, R., and Huerta-Sánchez, E. (2015). Evidence for archaic adaptive introgression in humans. Nature Reviews Genetics 2015 16:6, 16(6):359–371.

Reich, D., Patterson, N., Campbell, D., Tandon, A., Mazieres, S., Ray, N., Parra, M. V., Rojas, W., Duque, C., Mesa, N., García, L. F., Triana, O., Blair, S., Maestre, A., Dib, J. C., Bravi, C. M., Bailliet, G., Corach, D., Hünemeier, T., Bortolini, M. C., Salzano, F. M., Petzl-Erler, M. L., Acuña-Alonzo, V., Aguilar-Salinas, C., Canizales-Quinteros, S., Tusié-Luna, T., Riba, L., Rodríguez-Cruz, M., Lopez-Alarcón, M., Coral-Vazquez, R., Canto-Cetina, T., Silva-Zolezzi, I., Fernandez-Lopez, J. C., Contreras, A. V., Jimenez-Sanchez, G., Gómez-Vázquez, M. J., Molina, J., Carracedo, Á., Salas, A., Gallo, C., Poletti, G., Witonsky, D. B., Alkorta-Aranburu, G., Sukernik, R. I., Osipova, L., Fedorova, S. A., Vasquez, R., Villena, M., Moreau, C., Barrantes, R., Pauls, D., Excoffier, L., Bedoya, G., Rothhammer, F., Dugoujon, J. M., Larrouy, G., Klitz, W., Labuda, D., Kidd, J., Kidd, K., Di Rienzo, A., Freimer, N. B., Price, A. L., and Ruiz-Linares, A. (2012). Reconstructing Native American population history. Nature, 488(7411):370–374.

Reich, D., Thangaraj, K., Patterson, N., Price, A. L., and Singh, L. (2009). Reconstructing Indian population history. Nature, 461(7263):489–494.

Rheindt, F. E., Fujita, M. K., Wilton, P. R., and Edwards, S. V. (2014). Introgression and phenotypic assimilation in Zimmerius flycatchers (Tyrannidae): population genetic and phylogenetic inferences from genome–wide SNPs. Systematic biology, 63(2):134–152.

Rieseberg, L. H. and Wendel, J. F. (1993). lntrogression and Its Consequences in Plants. In Harrison, R. G., editor, Hybrid zones and the evolutionary process, chapter 4, pages 70–109. Oxford University Press.

Sankararaman, S. (2020). Methods for detecting introgressed archaic sequences. Current Opinion in Genetics & Development, 62:85–90.

Sankararaman, S., Mallick, S., Dannemann, M., Prüfer, K., Kelso, J., Pääbo, S., Patterson, N., and Reich, D. (2014). The genomic landscape of Neanderthal ancestry in present-day humans. Nature 2014 507:7492, 507(7492):354–357.

Seguin-Orlando, A., Korneliussen, T. S., Sikora, M., Malaspinas, A. S., Manica, A., Moltke, I., Albrechtsen, A., Ko, A., Margaryan, A., Moiseyev, V., Goebel, T., Westaway, M., Lambert, D., Khartanovich, V., Wall, J. D., Nigst, P. R., Foley, R. A., Lahr, M. M., Nielsen, R., Orlando, L., and Willerslev, E. (2014). Genomic structure in Europeans dating back at least 36,200 years. Science, 346(6213):1113– 1118.

Skov, L., Hui, R., Shchur, V., Hobolth, A., Scally, A., Schierup, M. H., and Durbin, R. (2018). Detecting archaic introgression using an unadmixed outgroup. PLOS Genetics, 14(9):e1007641.

Slatkin, M. (1985a). Gene Flow in Natural Populations. Source: Annual Review of Ecology and Systematics, 16:393–430.

Slatkin, M. (1985b). Rare alleles indicators of gene flow. Evolution; international journal of organic evolution, 39(1):53–65.

Slatkin, M. (1987). Gene Flow and the Geographic Structure of Natural Populations. Science, 236(4803):787–792.

Smith, J. and Kronforst, M. R. (2013). Do Heliconius butterfly species exchange mimicry alleles? Biology letters, 9(4).

Sousa, V. C., Fritz, M., Beaumont, M. A., and Chikhi, L. (2009). Approximate Bayesian Computation Without Summary Statistics: The Case of Admixture. Genetics, 181(4):1507–1519.

Steinrücken, M., Spence, J. P., Kamm, J. A., Wieczorek, E., and Song, Y. S. (2018). Model-based detection and analysis of introgressed Neanderthal ancestry in modern humans. Molecular Ecology, 27(19):3873–3888.

Suarez-Gonzalez, A., Lexer, C., and Cronk, Q. C. (2018). Adaptive introgression: a plant perspective. Biology Letters, 14(3).

Tang, H., Coram, M., Wang, P., Zhu, X., and Risch, N. (2006). Reconstructing Genetic Ancestry Blocks in Admixed Individuals. The American Journal of Human Genetics, 79(1):1–12.

Tournebize, R. and Chikhi, L. (2023). Questioning neanderthal admixture: on models, robustness and consensus in human evolution. bioRxiv.

Tung, J. and Barreiro, L. B. (2017). The contribution of admixture to primate evolution. Current Opinion in Genetics & Development, 47:61–68.

Vernot, B. and Akey, J. M. (2014). Resurrecting surviving Neandertal lineages from modern human genomes. Science, 343(6174):1017–1021.

Vernot, B., Tucci, S., Kelso, J., Schraiber, J. G., Wolf, A. B., Gittelman, R. M., Dannemann, M., Grote, S., McCoy, R. C., Norton, H., Scheinfeldt, L. B., Merriwether, D. A., Koki, G., Friedlaender, J. S., Wakefield, J., Pääbo, S., and Akey, J. M. (2016). Excavating Neandertal and Denisovan DNA from the genomes of Melanesian individuals. Science, 352(6282):235–239.

Wall, J. D., Lohmueller, K. E., and Plagnol, V. (2009). Detecting Ancient Admixture and Estimating Demographic Parameters in Multiple Human Populations. Molecular Biology and Evolution, 26(8):1823– 1827.

Wegmann, D. and Excoffier, L. (2010). Bayesian Inference of the Demographic History of Chimpanzees. Molecular Biology and Evolution, 27(6):1425–1435.

Wegmann, D., Kessner, D. E., Veeramah, K. R., Mathias, R. A., Nicolae, D. L., Yanek, L. R., Sun, Y. V., Torgerson, D. G., Rafaels, N., Mosley, T., Becker, L. C., Ruczinski, I., Beaty, T. H., Kardia, S. L., Meyers, D. A., Barnes, K. C., Becker, D. M., Freimer, N. B., and Novembre, J. (2011). Recombination rates in admixed individuals identified by ancestry-based inference. Nature Genetics 2011 43:9, 43(9):847–853.

Wu, C. I. (2001). The genic view of the process of speciation. Journal of Evolutionary Biology, 14(6):851– 865.

Yamamichi, M. and Innan, H. (2012). Estimating the migration rate from genetic variation data. Heredity, 108(4):362–363.

Yang, M. A., Malaspinas, A. S., Durand, E. Y., and Slatkin, M. (2012). Ancient structure in Africa unlikely to explain Neanderthal and non-African genetic similarity. Molecular biology and evolution, 29(10):2987–2995.

Zhang, X., Witt, K. E., Bañuelos, M. M., Ko, A., Yuan, K., Xu, S., Nielsen, R., and Huerta-Sanchez, E. (2021). The history and evolution of the Denisovan-EPAS1 haplotype in Tibetans. Proceedings of the National Academy of Sciences of the United States of America, 118(22):e2020803118.

