## Supplementary Material for "Inference of Locus-Specific Population Mixtures From Linked Genome-Wide Allele Frequencies"

#### Contents

**1 Inference** **S1**
1.1 Optimization of Emission Probabilities . . . . . S2
1.2 Optimization of Transition Probabilities . . . . . S3

**2 Initialization** **S4**
2.1 Step 1: A Gaussian Mixture Model . . . . . S4
2.1.1 Inference . . . . . S5
2.1.2 Initial guess and choice of number of classes . . . . . S6
2.2 Step 2: Guessing initial branch length . . . . . S6

**3 References** **S7**

### 1 Inference

We have to infer the transition and emission probabilities from the observations. Since we do not know through which hidden states the trajectories went, we apply the Baum-Welch algorithm to estimate the following parameters $\boldsymbol{\theta}$ of the HMM:

1. the initial state distribution $\eta_j = \mathbb{P}(z_1 = j)$;
2. the transition matrix parameters, where $i$ is associated to each migration edge:

    $\kappa_i$ (scaling factors),
    $\phi_i$ (control number of sites not in "neutral state"),
    $\zeta_i$ (control extent of sites not in "neutral state"),
    $a_i$ (attractors, "neutral states");

3. the branch lengths $\boldsymbol{c} = (c_1, \ldots, c_k)$;
4. the parameters $\mu, \sigma^2$ of the root state prior.

Applying the Baum-Welch algorithm to HMM is rather standard (Baum et al., 1970), but it involves lengthy calculations for this rather complex model, as we will show in the following. The Baum-Welch algorithm is a variant of the Expectation-Maximization (EM) algorithm which updates the parameters by alternating between an E-step and an M-step. Let $\boldsymbol{\theta}'$ denote the old and $\boldsymbol{\theta}$ the new parameters. In the E-step of the algorithm, we calculate the expected complete data log-likelihood given by

$$\begin{aligned} Q(\boldsymbol{\theta}, \boldsymbol{\theta}') = &\sum_j \gamma_1(j) \log \eta_j + \sum_l \sum_{j,j'} \gamma_l(j,j') \log \mathbb{P}_l(j,j') \\ &+ \sum_l \sum_j \gamma_l(j) \log \pi(\mathbf{d}_l | \mu, \sigma^2, \mathcal{G}_j), \end{aligned} \tag{S.1}$$

where we used the notation from the main text and where

$$\gamma_l(j) = \mathbb{P}(z_l = j \mid \mathbf{d}_{1:L}, \boldsymbol{\theta}')$$

and

$$\gamma_l(j,j') = \mathbb{P}(z_{l-1} = j, z_l = j' \mid \mathbf{d}_{1:L}, \boldsymbol{\theta}').$$

These $\gamma$'s can be calculated by the standard forward-backward algorithm for HMMs (Murphy, 2012). Maximization w.r. to the new parameters is standard for the initial state distribution: $\eta_j^* = \gamma_1(j)$. For each EM iteration, we also need to store all $\gamma$'s to calculate the derivatives shown below.

#### 1.1 Optimization of Emission Probabilities

To infer the branch lengths, we use the Newton-Raphson (NR) algorithm (Nocedal and Wright, 2006; Lange, 2010). To avoid boundary issues, we allow for negative branch lengths as long as they result in valid variance-covariance matrices $\boldsymbol{W}_j$. Specifically, we check that i) all diagonal entries of $\boldsymbol{W}_j$ are $\geq 0$ and that ii) the matrix is positive semi-definite. We do so for all boundary cases, i.e. for the $\boldsymbol{W}_j$ of all combinations with either zero or full migration on each migration edge (e.g. (0,0), (0,1), (1,0) and (1,1) in case of two migration edges). However, we output a warning whenever negative branch lengths are inferred as they are often a sign of a misspecified tree.

The gradient of $Q(\boldsymbol{\theta}, \boldsymbol{\theta}')$ w. r. to the branch lengths is

$$\frac{\partial}{\partial c_k} Q(\boldsymbol{\theta}, \boldsymbol{\theta}') = \sum_l \sum_j \gamma_l(j) \frac{\partial}{\partial c_k} \log \pi(\mathbf{d}_l | \mu, \sigma^2, \mathcal{G}_j). \tag{S.2}$$

The terms $\frac{\partial}{\partial c_k} \log \pi(\mathbf{d}_l | \mu, \sigma^2, \mathcal{G}_j)$ can be evaluated by defining the matrix $\boldsymbol{J}_{jk}$ given by (6) . (Note that there we did not yet have the index $j$ because at this stage the matrix was defined for the value $w$ instead of $w_j$.) From (7) we get:

$$\frac{\partial}{\partial c_k} \boldsymbol{S}_{lj} = \frac{\partial}{\partial c_k} \boldsymbol{W}_j = \boldsymbol{J}_{jk}.$$

We can now calculate the partial derivatives of all the terms in (15). The derivative of the identity $\boldsymbol{S}_{lj}^{-1} \boldsymbol{S}_{lj} = \boldsymbol{I}$ yields

$$\frac{\partial}{\partial c_k} \boldsymbol{S}_{lj}^{-1} = -\boldsymbol{S}_{lj}^{-1} \boldsymbol{J}_{jk} \boldsymbol{S}_{lj}^{-1}.$$

Using Jacobi's formula we get

$$\frac{\partial}{\partial c_k} \log \det \boldsymbol{S}_{lj} = \operatorname{tr}\left( \boldsymbol{S}_{lj}^{-1} \frac{\partial}{\partial c_k} \boldsymbol{S}_{lj} \right) = \operatorname{tr}\left( \boldsymbol{S}_{lj}^{-1} \boldsymbol{J}_{jk} \right).$$

With this information we obtain the partial derivatives

$$\begin{aligned} \frac{\partial}{\partial c_k} \log \pi(\mathbf{d}_l | \mu_l, \mathcal{G}_j) = &-\frac{1}{2} \operatorname{tr}\left( \boldsymbol{S}_{lj}^{-1} \boldsymbol{J}_{jk} \right) \\ &+ \frac{1}{2} (\mathbf{d}_l - \mu_l \mathbf{1})' \boldsymbol{S}_{lj}^{-1} \boldsymbol{J}_{jk} \boldsymbol{S}_{lj}^{-1} (\mathbf{d}_l - \mu_l \mathbf{1}) \end{aligned} \tag{S.3}$$

that form a gradient function of the branch lengths:

$$\boldsymbol{F}(\boldsymbol{c}) = \boldsymbol{F}(c_1, \ldots, c_K) := \left( \frac{\partial}{\partial c_1} Q(\boldsymbol{\theta}, \boldsymbol{\theta}'), \ldots, \frac{\partial}{\partial c_K} Q(\boldsymbol{\theta}, \boldsymbol{\theta}') \right)'.$$

As mentioned, we will use the NR algorithm to solve $\boldsymbol{F}(\boldsymbol{c}) = (0, \ldots, 0)'$, for which we also need the Jacobian of $\boldsymbol{F}$. From (eq. S.2) and (eq. S.3) we obtain

$$\frac{\partial}{\partial c_i}\boldsymbol{F}_k(\boldsymbol{c}) = \frac{1}{2}\sum_l \sum_j \gamma_l(j)\Big[\mathrm{tr}\boldsymbol{K}_{lj,ik} - (\mathbf{d}_l - \mu_l\mathbf{1})'\left(\boldsymbol{K}_{lj,ik} + \boldsymbol{K}_{lj,ki}\right)\boldsymbol{S}_{lj}^{-1}(\mathbf{d}_l - \mu_l\mathbf{1})\Big], \tag{S.4}$$

where we introduce the notation

$$\boldsymbol{K}_{lj,ik} = \boldsymbol{S}_{lj}^{-1}\boldsymbol{J}_{ji}\boldsymbol{S}_{lj}^{-1}\boldsymbol{J}_{jk}.$$

The parameter $\mu$ of the root state prior can be inferred explicitly if we know $\sigma^2$. We get

$$\mu* = \frac{\sum_l \sum_j \gamma_l(j)\mathbf{1}'\boldsymbol{S}_{lj}^{-1}\mathbf{d}_l}{\sum_l \sum_j \gamma_l(j)\mathbf{1}'\boldsymbol{S}_{lj}^{-1}\mathbf{1}} \tag{S.5}$$

where we have set $\boldsymbol{S}_{lj} = \boldsymbol{\Sigma}_l + \boldsymbol{W}_j + \sigma^2\mathbf{1}\mathbf{1}'$, see Eq. (14).

There is no simple formula for $\sigma^2$ and we optimize for it with the Newton-Raphson algorithm. To this end, we need the first and second derivatives of (eq. S.1) with respect to $\sigma^2$. Using Jacobi's formula, we obtain the expression

$$\frac{\partial}{\partial\sigma^2}Q(\boldsymbol{\theta}, \boldsymbol{\theta}') = -\frac{1}{2}\sum_{j,l}\gamma_l(j)\left[\mathrm{tr}(\boldsymbol{T}_{lj}) - (\mathbf{d}_l - \mu\mathbf{1})'\boldsymbol{T}_{lj}\boldsymbol{S}_{lj}^{-1}(\mathbf{d}_l - \mu\mathbf{1})\right],$$

wherein we write $\boldsymbol{T}_{lj} = \boldsymbol{S}_{lj}^{-1}\mathbf{1}\mathbf{1}'$. For the second derivative of $Q$ w.r. to $\sigma^2$ we get

$$\frac{\partial^2}{(\partial\sigma^2)^2}Q(\boldsymbol{\theta}, \boldsymbol{\theta}') = \frac{1}{2}\sum_{j,l}\gamma_l(j)\left[\mathrm{tr}(\boldsymbol{T}_{lj}^2) - 2(\mathbf{d}_l - \mu\mathbf{1})'\boldsymbol{T}_{lj}^2\boldsymbol{S}_{lj}^{-1}(\mathbf{d}_l - \mu\mathbf{1})\right].$$

#### 1.2 Optimization of Transition Probabilities

The proposed transition matrix is parametrized by $\kappa_i$, $\phi_i$ and $\zeta_i$, and additionally by an index $a_i$ that denotes the row index of the attractor (0 corresponding to the bottom row, 1 to the second row, and so on):

$$\boldsymbol{\Lambda}_i = \begin{pmatrix} -1 & 1 & 0 & 0 & \ldots & 0 & 0 & 0 & 0 \\ 1-\zeta_i & -2 & 1+\zeta_i & 0 & \ldots & 0 & 0 & 0 & 0 \\ 0 & 1-\zeta_i & -2 & 1+\zeta_i & \ldots & 0 & 0 & 0 & 0 \\ \vdots & \vdots & \vdots & \vdots & \ddots & \vdots & \vdots & \vdots & \vdots \\ 0 & 0 & 0 & 0 & \ldots & 1+\zeta_1 & -2 & 1-\zeta_i & 0 \\ 0 & 0 & 0 & 0 & \ldots & 0 & 1+\zeta_i & -2 & 1-\zeta_i \\ 0 & 0 & 0 & 0 & \ldots & 0 & 0 & 1 & -1 \end{pmatrix}$$

with the attractor row given by

$$\begin{pmatrix} 0 & \ldots & 0 & \phi_i & -2\phi_i & \phi_i & 0 & \ldots & 0 \end{pmatrix}.$$

The parameter $\zeta_i$ models the rate of moving away from the periphery towards the attractor. If $\zeta_i = 0$, it's equally likely to move up or down. If $\zeta_i = 1$, it is impossible to move to the periphery. The parameter $\phi_i$ models the rate of leaving the attractor. If $\phi_i = 0$, it's impossible to leave the attractor. If $\phi_i = 1$, the rate of leaving the attractor matches that of leaving any other state.

If there are only three states, the generating matrix reduces to:

$$\boldsymbol{\Lambda}_i = \begin{cases} \kappa_i \cdot \begin{pmatrix} -\phi_i & \phi_i & 0 \\ 1+\zeta_i & -2 & 1-\zeta_i \\ 0 & 1 & -1 \end{pmatrix} & \text{if } a_i = 0 \\ \kappa_i \cdot \begin{pmatrix} -1 & 1 & 0 \\ \phi_i & -2\phi_i & \phi_i \\ 0 & 1 & -1 \end{pmatrix} & \text{if } a_i = 1 \\ \kappa_i \cdot \begin{pmatrix} -1 & 1 & 0 \\ 1-\zeta_i & -2 & 1+\zeta_i \\ 0 & \phi_i & -\phi_i \end{pmatrix} & \text{if } a_i = 2. \end{cases}$$

Note that in the case of three states and $a_i = 1$ there is no $\zeta_i$.

The stationary is complicated to calculate explicitly and we obtain it by solving a system of normal equations. For optimizing the transition matrix parameters $\kappa_i$, $\phi$, $\zeta_i$ for a specific $a_i$, we use the Nelder-Mead (Nelder and Mead, 1965) algorithm.

### 2 Initialization

As the Baum-Welch algorithm is sensitive to good starting values, we developed a multi-step strategy to initialize the branch lengths $\boldsymbol{c}$, the root priors $\mu$ and $\sigma_2$ as well as the transition matrix parameters to reasonable values.

#### 2.1 Step 1: A Gaussian Mixture Model

In the first step, we seek to get a good estimate of the variance-covariance matrices $\boldsymbol{W}_l$ without imposing any constraints given by the graph $\mathcal{G}$. To account for possible variation in $\boldsymbol{W}_l$ across loci due to variation in migration strengths, we partition the loci into $R$ classes and infer the variance-covariance matrices $\mathbf{W}_r$ across populations for each class $r = 1, \ldots, R$.

We proceed as follows: We assume that the ancestral frequencies $\mu_l$, $l = 1, \ldots, L$, are drawn i.i.d. from the prior $\mathcal{N}(\mu_0, \sigma_0^2)$ and that the observations are the random vectors $\mathbf{d}_l$ that are drawn from a series of Gaussian mixture models (GMM) with density

$$\mathbf{d}_l \sim \sum_{r=1}^{R} \pi_r \mathcal{N}(\mathbf{d}_l | \mu_l \mathbf{1}, \mathbf{W}_r), \quad l = 1, \ldots, L,$$

where we use the abbreviation

$$\mathcal{N}(\mathbf{d}_l | \mu_l \mathbf{1}, \mathbf{W}_r) = \frac{1}{\sqrt{(2\pi)^M |\mathbf{W}_r|}} \exp\left[ -\frac{1}{2} (\mathbf{d}_l - \mu_l \mathbf{1})' \mathbf{W}_r^{-1} (\mathbf{d}_l - \mu_l \mathbf{1}) \right].$$

That is, the values $\mathbf{d}_l$ are drawn with probabilities (mixing weights) $\pi_r$ from one of the $R$ classes each of which produces random vectors from a multivariate normal distribution with mean $\mu_l \mathbf{1}$ and variance matrix $\mathbf{W}_r$. Clearly

$$\sum_{r=1}^{R} \pi_r = 1.$$

The parameters of this model are thus:

1. the variance matrices $\mathbf{W}_r$, $r = 1, \ldots, R$, for each of the $R$ classes of the GMM;
2. the mean $\mu_0$ and the variance $\sigma_0^2$ of the Gaussian prior for $\mu_l$;
3. the mixing weights $\pi_r$.

To simplify the notation we will call this whole set of parameters $\boldsymbol{\theta}$.

##### 2.1.1 Inference

We apply the EM algorithm in order to estimate these parameters. The hidden data for the EM algorithm are:

1. The mean vector $\boldsymbol{\mu} = (\mu_1, \ldots, \mu_L)$. These are continuous numbers that will be integrated out in the E-step.
2. The vector $\boldsymbol{s} = (s_1, \ldots, s_L)$ that indicates for each site which of the classes has been chosen.

**E-step.** The complete data likelihood is given by

$$L_c(\boldsymbol{\theta}; \mathbf{d}, \boldsymbol{\mu}, \boldsymbol{s}) = \prod_{l=1}^{L} \sum_{r=1}^{R} \mathrm{Ind}(r = s_l)\ \mathcal{N}(\mathbf{d}_l | \mu_l \mathbf{1}, \mathbf{W}_r)\ \frac{1}{\sqrt{2\pi}\sigma_0} e^{-\frac{1}{2\sigma_0^2}(\mu_l - \mu_0)^2}.$$

The expected complete data log-likelihood is calculated as

$$Q(\boldsymbol{\theta}; \boldsymbol{\theta}') = \mathbb{E}\left[\log L_c(\boldsymbol{\theta}; \mathbf{d}, \boldsymbol{\mu}, \boldsymbol{s}) | \boldsymbol{\theta}'\right]. \tag{S.6}$$

Using Bayes rule for linear Gaussian systems (see Murphy, 4.4.1), it is not hard to check that

$$\pi(\mu_l | \mathbf{d}_l, r, \boldsymbol{\theta}') = \mathcal{N}(\nu_{lr}, \rho_r^2)$$

where

$$\nu_{lr} = \frac{\mathbf{1}'\mathbf{W}_r'^{-1}\mathbf{d}_l + \mu_0'/\sigma_0'^2}{\mathbf{1}'\mathbf{W}_r'^{-1}\mathbf{1} + 1/\sigma_0'^2}, \quad \rho_r^2 = \frac{1}{\mathbf{1}'\mathbf{W}_r'^{-1}\mathbf{1} + 1/\sigma_0'^2}. \tag{S.7}$$

In order to calculate the expectation in (eq. S.6) we need the values $p_{lr}' = \mathbb{P}(r | \mathbf{d}_l, \boldsymbol{\theta}')$ which by Bayes' theorem are given by

$$p_{lr}' = \frac{\pi_r' \mathcal{N}\left(\mathbf{d}_l | r, \mu_0', \sigma_0'^2\right)}{\sum_{s=1}^{R} \pi_s' \mathcal{N}\left(\mathbf{d}_l | s, \mu_0', \sigma_0'^2\right)}$$

where

$$\mathcal{N}\left(\mathbf{d}_l | r, \mu_0', \sigma_0'^2\right) = \mathcal{N}\left(\mathbf{d}_l \mid \mu_0'\mathbf{1}, \mathbf{W}_r' + \sigma_0'^2 \mathbf{1}\mathbf{1}'\right).$$

**M-step.** First we maximize with respect to $\mathbf{W}_r$. Writing only the relevant part for the optimization, we get for the expected complete data log-likelihood

$$Q_1(\boldsymbol{\theta}; \boldsymbol{\theta}') = -\frac{1}{2} \sum_{l=1}^{L} \sum_{r=1}^{R} p_{lr}' \left( \log |\mathbf{W}_r| + \int_{-\infty}^{\infty} (\mathbf{d}_l - \mu_l \mathbf{1})' \mathbf{W}_r^{-1} (\mathbf{d}_l - \mu_l \mathbf{1})\ \pi(\mu_l | \mathbf{d}_l, r, \boldsymbol{\theta}')\ \mathrm{d}\mu_l \right).$$

Using the well-known law for the matrix derivative

$$\frac{\partial}{\partial \boldsymbol{W}^{-1}} \log |\boldsymbol{W}| = -\frac{\partial}{\partial \boldsymbol{W}^{-1}} \log |\boldsymbol{W}^{-1}| = -\boldsymbol{W},$$

we obtain

$$\frac{\partial}{\partial \mathbf{W}_r^{-1}} Q_1(\boldsymbol{\theta}'; \boldsymbol{\theta}') = \frac{1}{2} \sum_{l=1}^{L} p_{lr}' \left( \mathbf{W}_r - \int_{-\infty}^{\infty} (\mathbf{d}_l - \mu_l \mathbf{1})(\mathbf{d}_l - \mu_l \mathbf{1})'\ \pi(\mu_l | \mathbf{d}_l, r, \boldsymbol{\theta}')\ \mathrm{d}\mu_l \right).$$

Setting this expression equal to zero, we get the optimal value

$$\mathbf{W}_r^* = \frac{1}{p_r'} \sum_{l=1}^{L} p_{lr}' \int_{-\infty}^{\infty} (\mathbf{d}_l - \mu_l \mathbf{1})(\mathbf{d}_l - \mu_l \mathbf{1})'\ \pi(\mu_l | \mathbf{d}_l, r, \boldsymbol{\theta}')\ \mathrm{d}\mu_l.$$

where $p_r' = \sum_{l=1}^{L} p_{lr}'$. Using

$$\int_{-\infty}^{\infty} \mu_l\ \pi(\mu_l | \mathbf{d}_l, r, \boldsymbol{\theta}')\ \mathrm{d}\mu_l = \nu_{lr} \tag{S.8}$$

and

$$\int_{-\infty}^{\infty} \mu_l^2 \ \pi(\mu_l|\mathbf{d}_l, r, \boldsymbol{\theta}') \ \mathrm{d}\mu_l = \nu_{lr}^2 + \rho_r^2, \tag{S.9}$$

we get

$$\begin{aligned} \mathbf{W}_r^* &= \frac{1}{p'_r}\sum_{l=1}^{L} p'_{lr}\left[\mathbf{d}_l\mathbf{d}'_l - \nu_{lr}(\mathbf{1}\mathbf{d}'_l + \mathbf{d}_l\mathbf{1}') + (\nu_{lr}^2 + \rho_r^2)\mathbf{1}\mathbf{1}'\right] \\ &= \rho_r^2\mathbf{1}\mathbf{1}' + \frac{1}{p'_r}\sum_{l=1}^{L} p'_{lr}(\mathbf{d}_l - \nu_{lr}\mathbf{1})(\mathbf{d}_l - \nu_{lr}\mathbf{1})'. \end{aligned} \tag{S.10}$$

The relevant part for the optimization of the expected complete data log-likelihood (eq. S.6) with respect to $\mu_0, \sigma_0^2$ is

$$Q_2(\mu_0, \sigma_0^2; \boldsymbol{\theta}') = -\frac{1}{2}\sum_{l=1}^{L}\sum_{r=1}^{R} p'_{lr}\left(\log\sigma_0^2 + \frac{1}{\sigma_0^2}\int_{-\infty}^{\infty}(\mu_l - \mu_0)^2 \ \pi(\mu_l|\mathbf{d}_l, r, \boldsymbol{\theta}') \ \mathrm{d}\mu_l\right).$$

Thanks to (eq. S.8) and (eq. S.9) we can explicitly solve

$$\frac{\partial}{\partial\mu_0} Q_2(\mu_0, \sigma_0^2; \boldsymbol{\theta}') = 0$$

and we obtain

$$\mu_0^* = \frac{1}{L}\sum_{l=1}^{L}\sum_{r=1}^{R} p'_{lr}\nu_{lr}. \tag{S.11}$$

Similarly, we solve

$$\frac{\partial}{\partial\sigma_0^2} Q_2(\mu_0^*, \sigma_0^2; \boldsymbol{\theta}') = 0$$

and get

$$\sigma_0^{2*} = \frac{1}{L}\sum_{l=1}^{L}\sum_{r=1}^{R} p'_{lr}\left(\rho_r^2 + (\nu_{lr} - \mu_0^*)^2\right). \tag{S.12}$$

The new estimate for the mixing weights is

$$\pi_r^* = \frac{1}{L}\sum_{l=1}^{L} p'_{lr}. \tag{S.13}$$

##### 2.1.2 Initial guess and choice of number of classes

We use the observed variance-covariance matrix of the transformed observed frequencies as an initial guess of the variance covariance matrix $\boldsymbol{W}_0$ and slightly perturb it to get initial guess for the other classes.

As it is unclear how many classes to use, we start with a rather large number of lasses (usually $R = 5$) and then reduce the number of classes until the proportion of loci $\pi_r > \pi^*$ is larger than a pre-defined threshold $\pi^*$. We found that a relative large value of $\pi^* = 0.2$ gave best results.

#### 2.2 Step 2: Guessing initial branch length

The $\mathbf{W}_r$ inferred in the previous step are a function of two different components of our model, namely the variance-covariance given by the graph $\mathcal{G}$ through the branch lengths $\boldsymbol{c}$ and migration rates $\boldsymbol{w}$, as well as the variance that stems from binomial sampling of the data ($\boldsymbol{\Sigma}_l$). We therefore estimate the branch lengths $\boldsymbol{c}$ with the Nelder-Mead algorithm by minimizing the weighted Residuals Sums of Squares (RSS) between $\mathbf{W}_r$ and $\widehat{\boldsymbol{W}}_r = f(\boldsymbol{c})$ given by (7) of the main text, while accounting for $\boldsymbol{\Sigma}_l$. More precisely, let $\boldsymbol{c} = (c_1, \dots, c_K)'$, define the vectorizations

$$\hat{w} = \mathrm{vect}(\widehat{\boldsymbol{W}}), \quad \boldsymbol{j}_k = \mathrm{vect}(\boldsymbol{J}_k),$$

and build the matrix

$$\boldsymbol{J} = (\boldsymbol{j}_1 | \dots | \boldsymbol{j}_K) \,.$$

We can, for each $r$, recast (7) into the form

$$\widehat{w} = \boldsymbol{J}\boldsymbol{c} + \bar{\boldsymbol{s}}. \tag{S.14}$$

with

$$\bar{\boldsymbol{s}} = \text{vect}(\bar{\boldsymbol{\Sigma}})$$

where $\bar{\boldsymbol{s}}$ represents the vectorization of the weighted average of all $\boldsymbol{\Sigma}_l$ described in (10)

Before optimizing, the initial branch lengths $c_k$ can be estimated with standard OLS from (eq. S.14). To do so, we need an initial $\widehat{\boldsymbol{W}}$ and $\boldsymbol{J}$. We calculate the observed variance covariance matrix and use it as the initial $\widehat{\boldsymbol{W}}$ for all $r$, similar to the initial $\boldsymbol{W}$. To build the initial $\boldsymbol{J}$, we randomly sample migration weights $p$ from a uniform distribution or take values provided by the user. If any initial branch length is negative, it is set close to zero.

For optimization, we run Nelder-Mead to find the best combination of $\boldsymbol{c}$ and $p$ that results in the minimum weighted RSS

$$\overline{RSS} = \sum_{r=1}^{R} \sum (\boldsymbol{W}_r - \widehat{\boldsymbol{W}}_r)^2 * \pi_r \tag{S.15}$$

The Nelder-Mead operates on $\log(c)$ and $\text{logit}(p)$ space. If any proposed value is zero, a minimum value is reassigned before the log or logit transformation, similarly a maximum value other than one is reassigned before the logit transformation. To avoid issues with local minima, we run the Nelder-Mead a thousand times with different starting locations, and choose the branch length that result in the smallest RSS value.

### 3 References


Baum, L. E., Petrie, T., Soules, G., and Weiss, N. (1970). A Maximization Technique Occurring in the Statistical Analysis of Probabilistic Functions of Markov Chains. *The Annals of Mathematical Statistics*, 41(1):164–171.

Lange, K. (2010). *Numerical Analysis for Statisticians.* Statistics and Computing. Springer New York, New York, NY.

Murphy, K. P. (2012). *Machine learning: a probabilistic perspective.*

Nelder, J. A. and Mead, R. (1965). A simplex method for function minimization. *The Computer Journal*, 7(4):308–313.

Nocedal, J. and Wright, S. (2006). *Numerical Optimization.* Springer Science & Business Media.
